# UNC-16/JIP3 negatively regulates actin dynamics dependent on DLK-1 and microtubule dynamics independent of DLK-1 in regenerating neurons

**DOI:** 10.1101/484683

**Authors:** Sucheta S. Kulkarni, Vidur Sabharwal, Seema Sheoran, Atrayee Basu, Kunihiro Matsumoto, Naoki Hisamoto, Anindya Ghosh-Roy, Sandhya P. Koushika

## Abstract

Neuronal regeneration after injury depends on the intrinsic growth potential of neurons. UNC-16, a *C. elegans* JIP3 homologue, inhibits axonal regeneration by regulating regrowth initiation and rate of regrowth. UNC-16/JIP3 inhibits the regeneration promoting activity of DLK-1 long but acts additively to and independently of inhibitory DLK-1 short isoform. UNC-16/JIP3 promotes DLK-1 punctate localization in a concentration dependent manner limiting DLK-1 long availability at the cut site minutes after injury. UNC-16 negatively regulates actin dynamics dependent on DLK-1 and microtubule dynamics independent of DLK-1. The faster regeneration seen in *unc-16* does not lead to functional recovery. We propose a model where UNC-16/JIP3 plays its inhibitory role through tight temporal and spatial control of DLK-1 function. The dual inhibitory control by both UNC-16 and DLK-1 short calibrate the intrinsic growth promoting function of DLK-1 long in vivo.

## Introduction

Neurons in the adult nervous system have a limited ability to regenerate after injury. Studies have shown that the extent of neuronal regeneration depends on the balance between factors that inhibit and promote neuronal regeneration which can be both extrinsic and intrinsic to the injured neuron. Some extrinsic inhibitory factors in the adult central nervous system are Nogo and its receptor (Huo et al., 2013; Peng, Zhou, Hu, Fink, & Mata, 2010; Simonen et al., 2003) and Myelin associated glycoprotein (Domeniconi et al., 2002; Liu, Fournier, GrandPre, & Strittmatter, 2002) while glial cell line-derived neurotrophic factor and brain-derived neurotrophic factor (Boyd & Gordon, 2003) are extrinsic growth promoting factors. Likewise Oligodendrocyte myelin glycoprotein (Oertle et al., 2003; K. C. Wang et al., 2002) and p75 Neurotrophin Receptor (Yamashita, Higuchi, & Tohyama, 2002) are intrinsic growth inhibitory factors and factors like cAMP, cytokines, and growth cone actin cytoskeleton modulators like GAP-43 and CAP-23 are intrinsic neuronal growth promoters. (Bomze, Bulsara, Iskandar, Caroni, & Skene, 2001; Cai, Shen, De Bellard, Tang, & Filbin, 1999; Nikulina, Tidwell, Dai, Bregman, & Filbin, 2004; Qiu et al., 2002).

Another important factor downstream of the signalling cascade is the dynamics of cytoskeletal elements (Blanquie & Bradke, 2018; Chisholm, 2013; Hur, Saijilafu, & Zhou, 2012). Microtubule dynamics has been shown to be elevated and under the control of the early signalling cascade post-injury to neuron (Lu, Lakonishok, & Gelfand, 2015). It has been shown that regulation of microtubule dynamics and polarity in response to JNK signalling is crucial to initiate regeneration of an axon post-injury (Stone, Nguyen, Tao, Allender, & Rolls, 2010). The regenerating neurons also form new growth cones which are rich in actin, actin filaments and their associated proteins and are together responsible for both formation and motility of the growth cone (Gomez & Letourneau, 2014).

The balance between growth promoting and inhibiting factors determine the extent to which a neuron regenerates. This can be achieved in multiple ways, for instance several kinases have been reported to regulate the intrinsic growth potential of regenerating neurons such as, Protein Kinase C (Lee et al., 2013), S6 Kinase (Hubert, Wu, Chisholm, & Jin, 2014), MAP Kinases such as Erk (Hanz et al., 2003), JNK (Lindwall & Kanje, 2005) and Dual Leucine zipper Kinase (DLK) (Hammarlund, Nix, Hauth, Jorgensen, & Bastiani, 2009).

One means to regulate the activity of such kinases in a spatio-temporal manner is through scaffolding molecules. Such molecules can bring together several proteins serving to modulate entire signalling cascades. JIPs are a family of classical scaffolding molecules and they are known to be able to switch their roles from growth promoting to growth inhibiting based on their levels and states of activation or deactivation (Whitmarsh, 2006).

JIP3, a JNK interacting protein (JIP) family member, assists in the formation of a functional JNK signalling module by assembling specific components of the MAPK cascade which is known to modulate neuronal growth in various systems (Byrd et al., 2001; Ito et al., 1999; Kelkar, Gupta, Dickens, & Davis, 2000). JIP3 interacts with the specific kinases of the JNK signalling pathway i.e. JNK, MKK7 and MLK3. MAPKKKs are the most upstream kinases in the MAPK cascade and play a role in regulating signal specificity (Kyriakis & Avruch, 2012). DLK, a MAPKKK, regulates synapse formation and is essential for neuronal regeneration (Hammarlund et al., 2009; Nakata et al., 2005). DLK-1 activation is tightly regulated and occurs only in the presence of specific stimuli like neuronal injury (Nakata et al., 2005; Yan & Jin, 2012). Homodimerisation of DLK-1 is sufficient for its activation (Nihalani, Merritt, & Holzman, 2000) and JIP1, another JIP family scaffold protein, maintains DLK in an inactive state in part by preventing its dimerization (Nihalani, Wong, & Holzman, 2003). Thus, the interaction of scaffolding molecules and kinases they scaffold can regulate activity of the kinases involved in neuronal regeneration.

Axonal injury breaks the connection between the neuronal cell body and its postsynaptic targets disrupting its functions and potentially the behaviours that depend on the neuron (He & Jin, 2016). Multiple studies in various systems have demonstrated that behavioural recovery is possible post-injury if the injured neuron regrows and reconnects to form a functional circuit. In the mouse model, removing growth inhibitory extracellular matrix molecules like chondroitin sulphate proteoglycans (CSPGs) from the glial scar and blocking growth inhibitory Nogo-66 receptors has been shown to improve behavioural recovery in corticospinal axons post-injury (Bradbury et al., 2002; J. E. Kim, Liu, Park, & Strittmatter, 2004). Zebrafish have been reported to grow brainstem/spinal cord transition zone axons post-transection and recover swimming behaviour (Becker, Wullimann, Becker, Bernhardt, & Schachner, 1997; Briona & Dorsky, 2014). In *C. elegans* there are multiple reports of partial recovery of locomotion following axotomy in multiple motor neurons (Byrne et al., 2016; El Bejjani & Hammarlund, 2012; Neumann, Nguyen, Hall, Ben-Yakar, & Hilliard, 2011; Yanik et al., 2004). In *C. elegans* it has been specifically shown that there is axon fusion followed by functional recovery in the touch neurons (Abay et al., 2017; Basu et al., 2017). These studies establish a link between axonal fusion and restoration of the functional neuronal circuit leading to recovery of the lost behavioural response.

To assess the role of scaffolding molecules in axon regeneration we investigated the role of a well-established JIP family protein in regeneration. We show that UNC-16, the *C. elegans* JIP3 homologue (Byrd et al., 2001), inhibits regeneration of touch neurons after axotomy. We show that the faster regeneration in *unc-16* animals is a result of quicker initiation of neuronal regrowth and is independent of Kinesin-1, Dynein and JNK-1 but is dependent on DLK-1. Our data suggest that UNC-16 independently and in a concentration dependent manner inhibits regeneration by acting on the regeneration promoting long isoform of DLK-1. UNC-16’s inhibitory effect on DLK-1 long is independent of and in addition to the inhibition provided by DLK-1 short. Our data suggest that UNC-16’s inhibitory activity likely occurs by sequestering DLK-1 long in a concentration dependent manner and preventing DLK-1 long accumulation at injury site. UNC-16 negatively regulates actin dynamics dependent on DLK-1 and microtubule dynamics independent of DLK-1. We also show that the faster regeneration seen in loss of UNC-16 does not translate into functional recovery, but slower regeneration observed when UNC-16 is over-expressed shows functional recovery comparable to wild type animals. We propose a model where UNC-16/JIP3 plays its regulatory role through tight temporal and spatial control of DLK-1 function. The dual inhibitory control by both UNC-16 and DLK-1 short can calibrate the intrinsic growth promoting function of DLK-1 long *in vivo* and a slower regeneration rate through modulation of cytoskeletal dynamics may assist in functional recovery of the injured neuron.

## Results

### Loss of UNC-16 leads to an increase in neuronal regeneration

To study UNC-16’s role in regeneration, we compared the post-axotomy regrowth in the posterior lateral microtubule (PLM) neuron of *unc-16* and wild type animals. We observed regeneration in the *ju146, e109, tb109*, *n730* and *km75* alleles of *unc-16*, with mutations at different locations (Fig. 1A) (Byrd et al., 2001; Choudhary et al., 2017; Edwards et al., 2013). *tb109*, *n730* and *km75* have stop codons which are predicted to truncate the protein at different lengths with *tb109* resulting in the shortest protein (Fig. 1A). Besides the known mutations in the protein, we also assessed and found reduced *unc-16* RNA levels in four of the five above alleles compared to wild type (Sup. Fig. 1). Our experiments were carried out 9 hours post-axotomy as they gave us clear differences between genotypes, data are represented as percentage regeneration that refers to the number of animals that show neuronal regeneration. All five alleles showed greater number of animals whose neurons regenerate as compared to wild type (Fig. 1A). The inhibitory effect of *unc-16* on neuronal regeneration is not allele specific varying from 43% to 88% in different loss of function (lf) alleles. In genetic screens for *C. elegans* motor neuron regeneration in *unc-70/β-spectrin* mutants an inhibitory role for *unc-16* has been reported (Chen et al., 2011; Nix et al., 2014).

**Figure 1.**
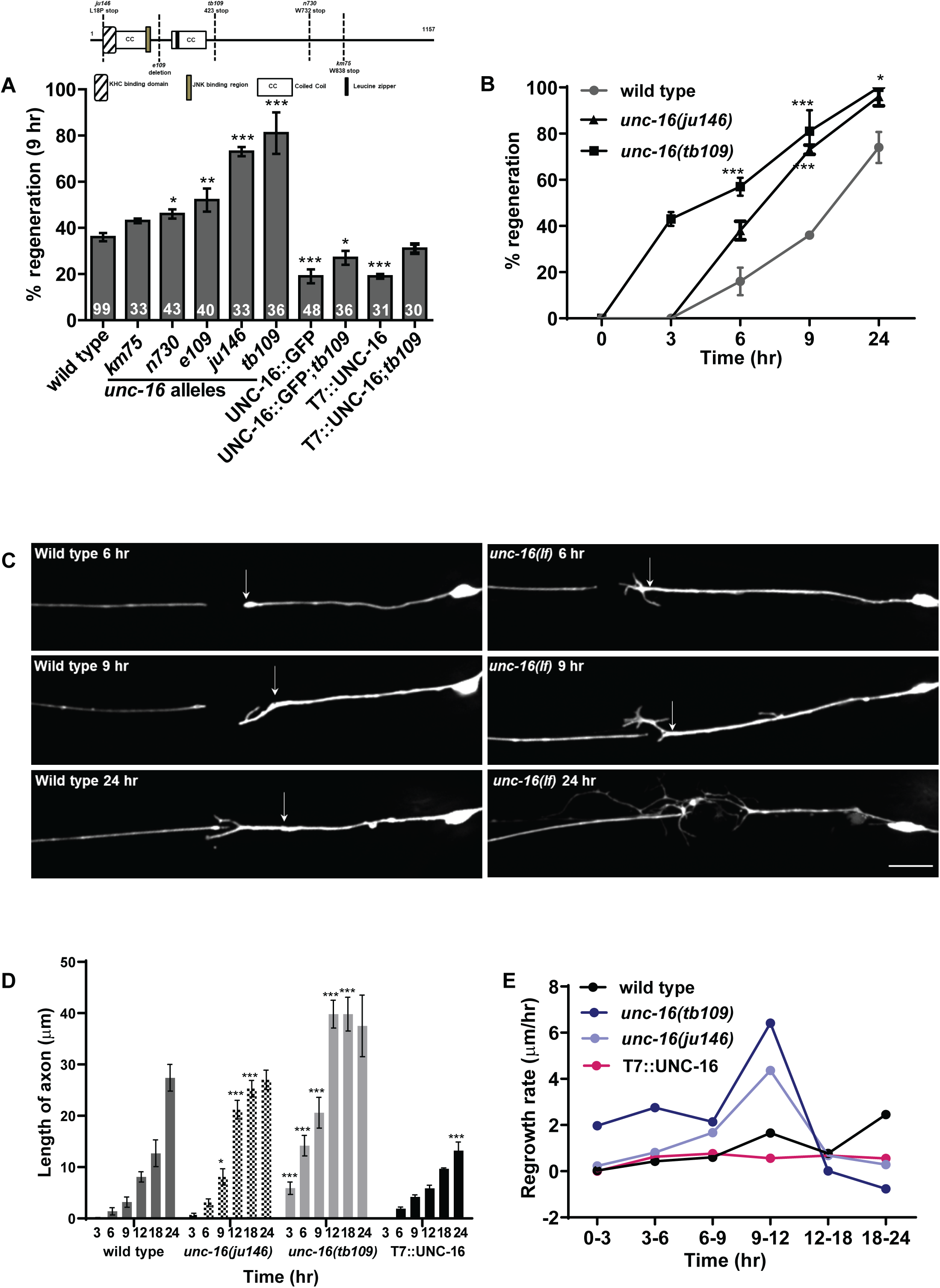
Loss of UNC-16/JIP3 promotes neuronal regeneration. ***A***, Percentage neurons regenerating post-axotomy in *unc-16* alleles, after expression of UNC-16::GFP and T7::UNC-16 in wild type and *unc-16(tb109)*. Inset: Schematic of UNC-16 domain and mutations. ***B***, Percentage neurons regenerating at specified time-points post-axotomy in wild type, *unc-16(ju146)* and *unc-16(tb109)*. ***C***, Representative images of the PLM process at specified time-points post-axotomy. Arrow indicates position of axotomy. Scale bar 10 µm. ***D***, Length of the regenerated neuronal process at different time points after axotomy. *E*, Regrowth rate of the neuronal process in each three-hour interval after axotomy. One-way ANOVA, p-value *<0.05, **<0.005, ***<0.001. Number of animals inside bars in *A*. Number of animals is 20 for *B*, *D* and *E*.

We found that expression of transgene encoded UNC-16::GFP (Byrd et al., 2001) or T7::UNC-16 in *unc-16(tb109)* animals reduces regeneration from 81% to 27% (Fig. 1A). Further, expression of UNC-16::GFP or T7::GFP in wild type animals resulted in 19% of the severed processes regenerating compared to 36% in wild type. This suggests that the presence and levels of UNC-16 is sufficient to inhibit regeneration following injury.

### Loss of UNC-16 results in faster initiation of regeneration

To examine the time course of regeneration in *unc-16*, we monitored the neuronal processes of *unc-16(tb109)* and *unc-16(ju146)* at different time points after axotomy. At 6 hours, 38% of *unc-16(ju146)* and 57% of *unc-16(tb109)* animals initiated neuronal regrowth compared to only 16% in wild type (Fig. 1B, C). Furthermore, neuronal outgrowth in both alleles of *unc-16* examined started earlier at 3 hours post axotomy unlike in wild type injured axons where it began at 6 hours (Fig. 1B, D). At 3 hrs post-axotomy, *unc-16(tb109)* neurons were 5.90 µm long whereas the wild type neuron does not show any growth. At 9 hrs *unc-16(ju146)* and *unc-16(tb109)* show a ∼2.6 and ∼6.7 fold significantly greater neuronal process length compared to wild type (Fig. 1B). The increase in the number of animals showing regrowth and the greater length of neuronal outgrowth together show that loss of *unc-16* promotes earlier initiation of regrowth. This increased number of animals that show neuronal regrowth persists even 24 hours post-axotomy (Fig. 1B). Over-expression of UNC-16 in neurons did not alter the time when regrowth was initiated, where growth began similar to wild type animals at 6 hours post-axotomy (Fig. 1D).

We also examined the rate of regrowth after initiation by measuring the length of the regrowing axon at defined times after axotomy. 6-9 hours post-axotomy, the neuronal processes in *unc-16(tb109)* animals grew at the rate of 2.1 µm/hour, significantly higher than the wild type neuronal growth rate of 0.6 µm/hour (Fig. 1E). This led to a six-fold increase in the regenerated process length in *unc-16(tb109)* compared to wild type animals 9 hours post-axotomy (Fig. 1D). The difference in growth rate between wild type (1.7 µm/hour) and *unc-16(tb109)* (6.4 µm/hour) animals was greatest in the 9-12 hour time interval post-axotomy. This faster growth rate led to a five-fold increase in neuronal length 12 hours post-axotomy in *unc-16(tb109)* compared to wild type. We also evaluated the growth rate in *unc-16(ju146)* and found similar trends (Fig 1E). At the 6-9 hour post-axotomy, neuronal processes in *unc-16(ju146)* grew at the rate of 1.7 µm/hour (Fig. 1E). They exhibit the highest growth rate of 4.4 µm/hour in the 9-12 hour time interval that reflects in the 2.5-fold increase in neuronal length 12 hours post-axotomy in *unc-16(ju146)* compared to wild type. Elevating UNC-16 levels using a transgene expressing T7:: UNC-16 reduces the growth rate (Fig. 1E) eventually resulting in shorter process lengths 24 hours post-axotomy (Fig. 1D).

At 18 hours post-axotomy the neuronal regrowth rate in *unc-16(tb109)* reduced and became comparable to wild type (Fig. 1E). 24 hours post-axotomy, more than 50% of the processes of *unc-16(tb109)* show several branches (Fig. 1C), some that overlap with the distal process, thus an accurate assessment of true growth rates is not readily measurable. Taken together our data suggest that the loss of *unc-16* results in a quicker initiation of regrowth and a faster rate of process regrowth up to 12 hours post-axotomy.

Our findings show that UNC-16 acts as an inhibitory factor in the early steps of axon regeneration and prevents regrowth initiation and slows the rate of axon extension after injury.

### Increased regeneration in *unc-16* depends on DLK-1

UNC-16 is a scaffolding protein that plays multiple roles in intracellular transport and MAPK signalling. It interacts with the Kinesin-1 motor (Sun, Zhu, Dixit, & Cavalli, 2011), the Dynein motor (Arimoto et al., 2011), a RUN domain protein UNC-14 (Sakamoto et al., 2005), a leucine-rich repeat kinase 2 (LRRK2) (Choudhary et al., 2017; Hsu, Chan, & Wolozin, 2010) and MAP kinases such as JNK-1 (Kelkar et al., 2000) and DLK (Ghosh et al., 2011). We thus examined if UNC-16 dependent regeneration depended on the above proteins.

We assessed neuronal regeneration in loss-of-function (lf) mutants of the Kinesin heavy chain-1 (*unc-116*), the JNK MAP kinase (*jnk-1*), RUN domain protein (*unc-14*), leucine-rich repeat kinase 2 (*lrk-1*), Dynein heavy chain (*dhc-1*) and the Dual-leucine zipper kinase (*dlk-1*). The single mutants either regenerate similar to wild type as seen in the *unc-116* mutant and the *jnk-1* mutant or regenerate significantly lower as seen in mutants of *unc-14*, *lrk-1*, *dhc-1* and *dlk-1* (Fig. 2A, Sup. Fig. 2 and 3). In addition, double mutants between *unc-16(tb109)* and the above-mentioned genes did not influence UNC-16 dependent regeneration (Fig 2A, Sup. Fig. 2 and 3) with the exception of *dlk-1*. We examined the effect of loss of UNC-16 in different alleles of *dlk-1* (Fig. 2A, Sup. Fig. 4). The milder *dlk-1* alleles *ju477* and *ju591* (Yan & Jin, 2012) (Sup. Fig. 4) do not affect regeneration on their own but they inhibit regeneration in *unc-16(tb109).* Regeneration in *dlk-1; unc-16(lf)* double mutants built with stronger alleles of *dlk-1*, viz. *ju476* and *tm4024*, is completely abolished (Fig. 2A). Thus the inhibitory role of UNC-16 in neuronal regeneration is dependent on DLK-1.

**Figure 2.**
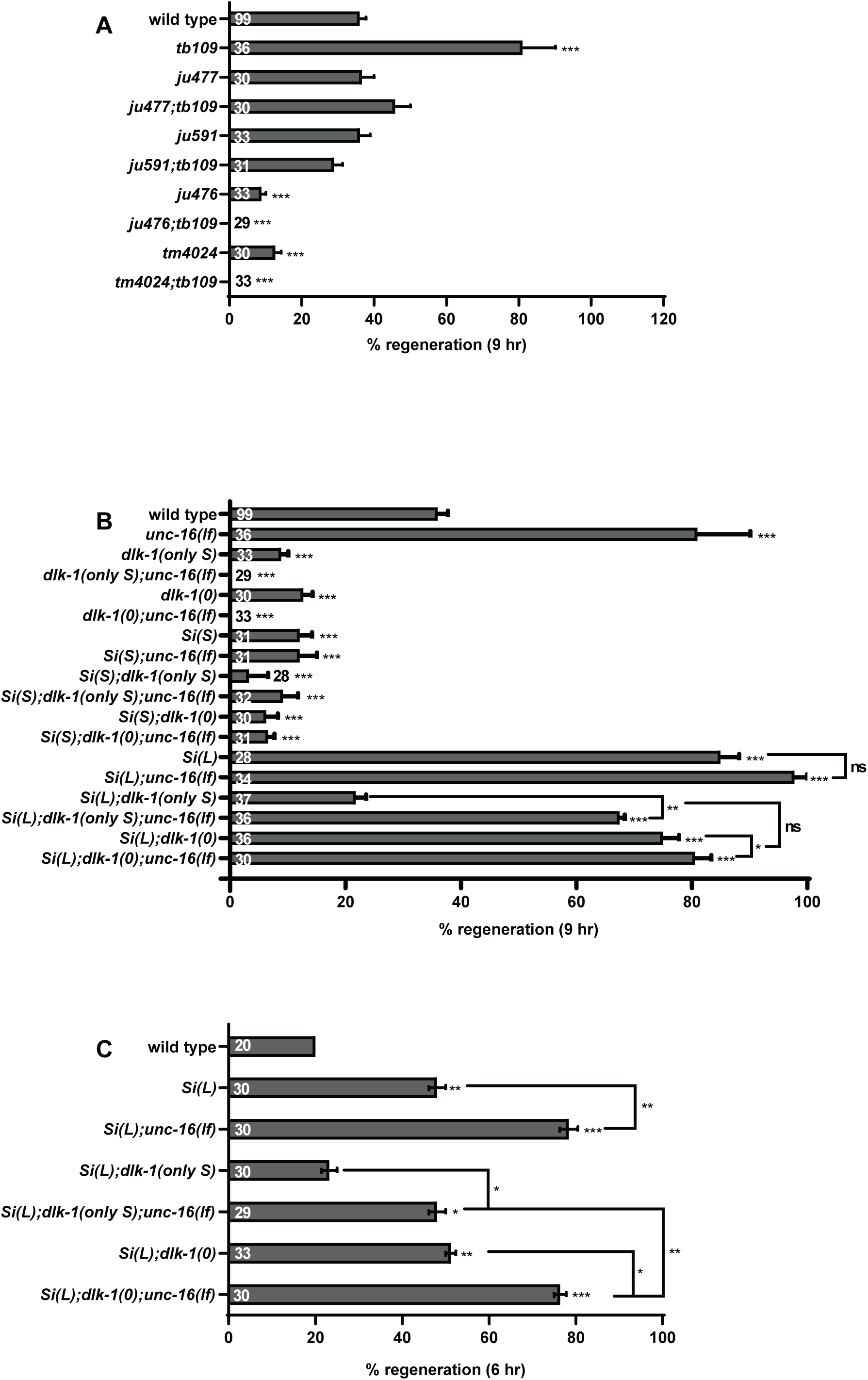
UNC-16 inhibits DLK-1L. ***A***, Effect of *unc-16* on regeneration phenotype of various alleles of *dlk-1*. ***B***, Genetic analysis of effect of *unc-16* in single copy DLK-1L and DLK-1S expressing strains (*MosSCI*) in wild type and *dlk-1* background 9 hours post-axotomy. ns: not significant. ***C***, Genetic analysis of effect of *unc-16(tb109)* in single copy DLK-1L in wild type and *dlk-1* background 6 hours post-axotomy. One-way ANOVA, p-value *<0.05, **<0.005, ***<0.001. Number of animals inside bars.

**Figure 3.**
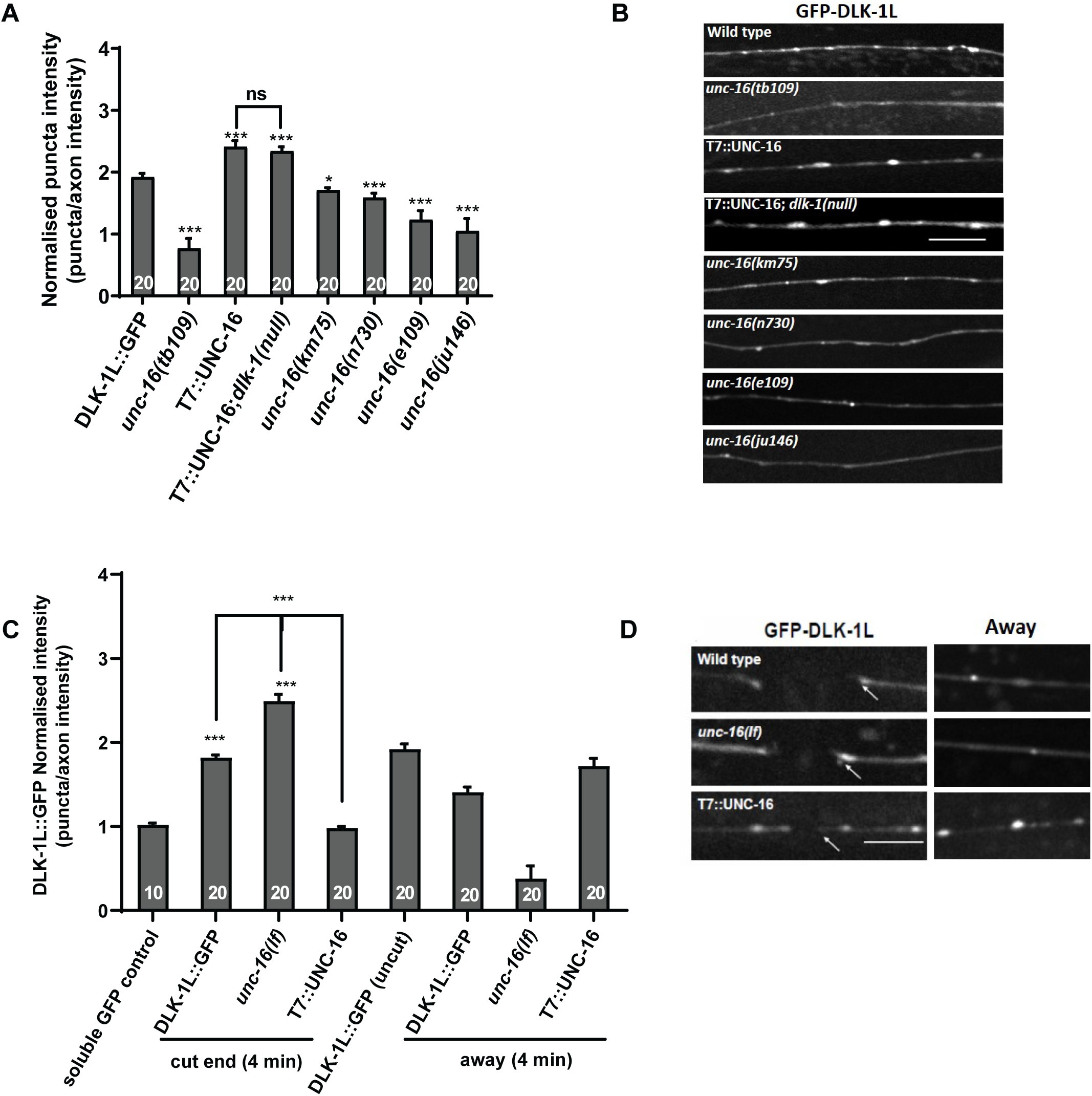
UNC-16 regulates DLK-1L in a dose-dependent fashion. ***A***, Normalised intensity quantification of effect of *unc-16(tb109)* and T7::UNC-16 on GFP::DLK-1L localization and in different alleles of *unc-16*. Intensity of *mec-4p::mCherry* was used as a negative soluble GFP control. ***B***, Representative pictures of GFP::DLK-1L localisation quantified in ***A***. Scale bar 10 µm. ***C***, Normalised intensity quantification of GFP::DLK-1L in the first 3 µm near the proximal cut end of an axotomised PLM neuron and away. ***D***, Representative picture of axotomised PLM neurons expressing GFP::DLK-1L, 4 minutes post-axotomy at the site of axotomy and away. Arrow points to the proximal cut end with respect to cell body. Scale bar 5 µm. One-way ANOVA, p-value *<0.05, ***<0.001. Number of animals inside bars.

**Figure 4.**
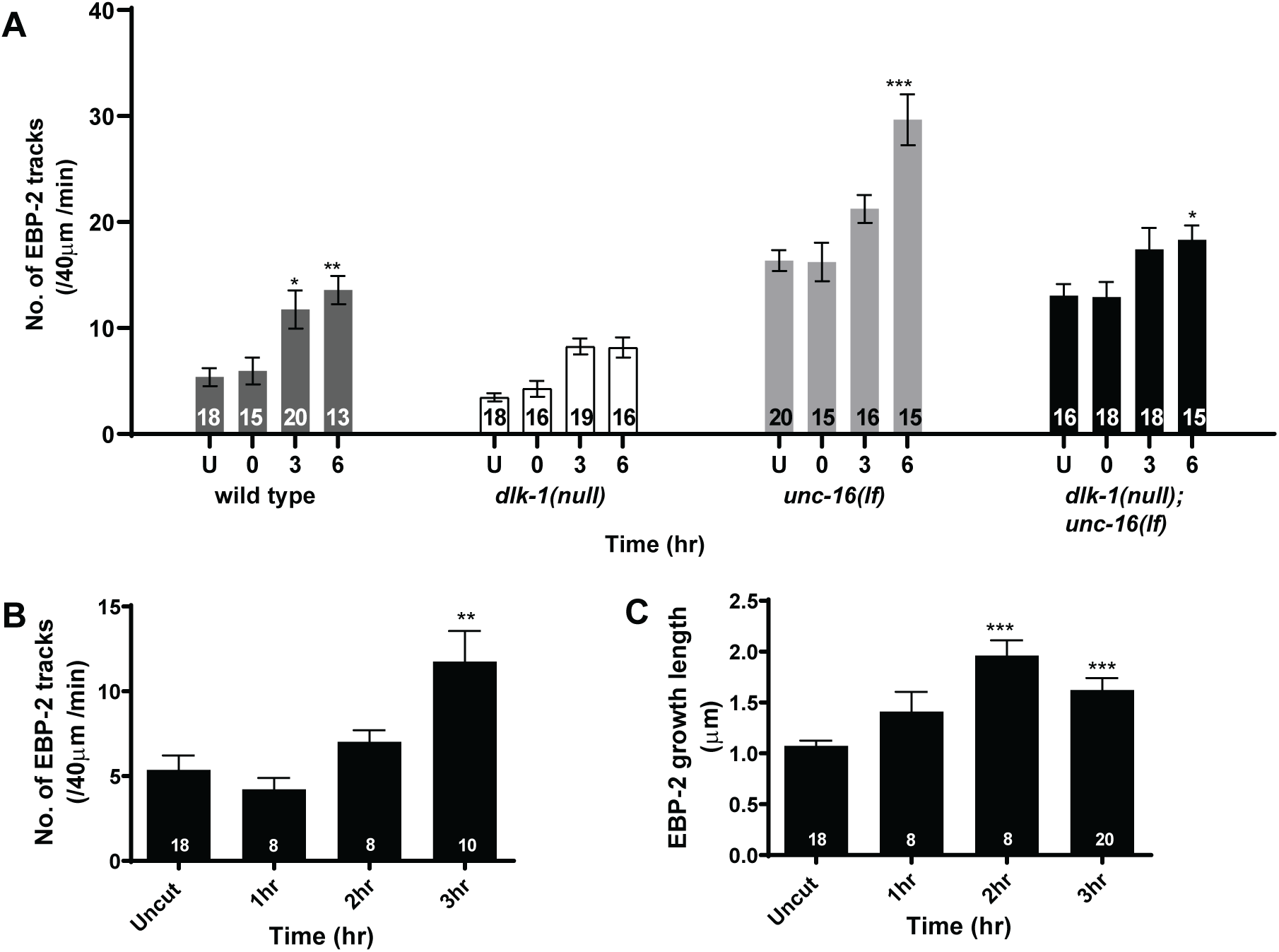
UNC-16 inhibits microtubule dynamics. ***A***. Numbers of EBP-2::GFP tracks in kymographs before and after axotomy in wild type and mutants in uncut and other timepoints as specified post-axotomy. ***B***. Number of EBP-2::GFP tracks in uncut and axotomized wild type animals as a function of time up to 3 hr post injury. ***C***. Change in length of EBP-2::GFP tracks in uncut and axotomized wild type animals over 3 hr post injury period. Number of animals inside bars. One-way ANOVA p-value *<0.05, **<0.005, ***<0.001.

### UNC-16 inhibits the regeneration promoting function of DLK-1L

DLK is an essential MAPKKK that functions in the initiation of regrowth of neuronal processes after injury and in growth cone formation in many model systems (Hammarlund et al., 2009; Shin et al., 2012; Z. Wang & Jin, 2011; Xiong et al., 2010; Yan, Wu, Chisholm, & Jin, 2009). *dlk-1* in *C. elegans* encodes two isoforms, DLK-1L (long) and DLK-1S (short), that have antagonistic functions. DLK-1S forms a heterodimer with DLK-1L and inactivates the regeneration promoting role of DLK-1L (Yan & Jin, 2012) (Sup. Fig. 4). JIP3, the mammalian orthologue of UNC-16, is known to form a complex with DLK (Holland et al., 2016; Kelkar et al., 2000). Hence we determined whether UNC-16 influences neuronal regeneration through a specific isoform of DLK-1.

We assessed the effect of loss of UNC-16 on neuronal regeneration in animals expressing a single copy of DLK-1S or DLK-1L. Consistent with previous studies, expression of a single copy of DLK-1S or DLK-1L in wild type respectively suppresses and enhances neuronal regeneration (Fig. 2B) (Yan & Jin, 2012). Similarly, results consistent with past studies were observed when either *dlk-1(ju476)* (does not express DLK-1L but expresses endogenous levels of intact DLK-1S) referred to as *dlk-1(onlyS)* or *dlk-1(tm4024)* (a DLK-1 null) referred to as *dlk-1(0)*, expressed either a single copy of DLK-1S or DLK-1L (Yan & Jin, 2012).

To determine if DLK-1S and UNC-16 act independently to inhibit DLK-1L mediated regeneration we assessed regeneration in the following genotypes: Single copy DLK-1L with *dlk-1(onlyS); unc-16(lf)* and *dlk-1(0); unc-16(lf)*. Single copy DLK-1L is introduced so that some regeneration occurs allowing us to assess the inhibitory effects of DLK-1S and UNC-16. We see that *unc-16(lf)* mutants lacking DLK-1S show significantly higher regeneration compared to animals that contain DLK-1S suggesting that DLK-1S and UNC-16 can both inhibit DLK-1L (Fig. 2B).

Loss of *unc-16* is unable to promote regeneration in animals expressing DLK-1S in wild type, *dlk-1(onlyS)* or *dlk-1(0)* backgrounds (Fig. 2B). However, animals expressing DLK-1L in *dlk-1(onlyS); unc-16(lf)* had enhanced regeneration compared with animals expressing DLK-1L in *dlk-1(onlyS)* alone (Fig. 2B). Additionally, animals expressing DLK-1L in *dlk-1(0); unc-16(lf)* further enhanced regeneration compared with animals expressing DLK-1L in *dlk-1(0)* greatest at 6 hours post-axotomy consistent with UNC-16’s role early in regeneration (Fig. 2B). Moreover, the loss of UNC-16 leads to a two-fold increase in regeneration in *dlk-1(onlyS)* animals and a further 1.5-fold increase in regeneration in *dlk-1(0)* (Fig. 2B). However, in *dlk-1(0)*, 9 hours post-axotomy, these differences were not very significant (Fig. 2C) and this may be due to the larger amounts of DLK-1L available in the single copy DLK-1L line, soon after axotomy, resulting in faster initiation of regeneration. Our data shows that UNC-16 inhibits the growth promoting activity of DLK-1L. We think UNC-16 acts independently and in addition to DLK-1S to inhibit regeneration since both *unc-16(lf)* and lack of DLK-1S enhance each other (Fig. 2B, C) and the inhibition by overexpression of DLK-1S is not suppressed by the *unc-16(lf)* mutation (Fig. 2C).

### UNC-16 changes the localization of DLK-1L

Vertebrate DLK, *C. elegans* GFP::DLK-1L, GFP::DLK-1S and endogenous UNC-16 are all shown to localize in punctae along the neuron (Edwards et al., 2013; Holland et al., 2016; Yan & Jin, 2012). The punctate localisation of vertebrate DLK has been shown to depend on palmitoylation dependent binding of DLK to JIP3 (Holland et al., 2016). Therefore, we examined the localization of DLK-1L in both *unc-16(tb109)* and in neurons with elevated levels of UNC-16.

The punctate localization of GFP::DLK-1L is lost in all six *unc-16* alleles examined (Fig. 3A, B, Sup Fig. 5). Further, upon UNC-16 over-expression, GFP::DLK-1L showed a significant increase in the intensity of puncta in wild type and in *dlk-1(0)* animals. This indicates that there is increased GFP::DLK-1L in each puncta in the presence of excess UNC-16, however the density of the puncta remain unaffected at ∼2.5±0.4/10 µm remaining similar to wild type (Fig. 3A, B). Our observations show that the punctate DLK-1L localization is dependent on the presence and levels of UNC-16 (Fig. 3A, B).

**Figure 5.**
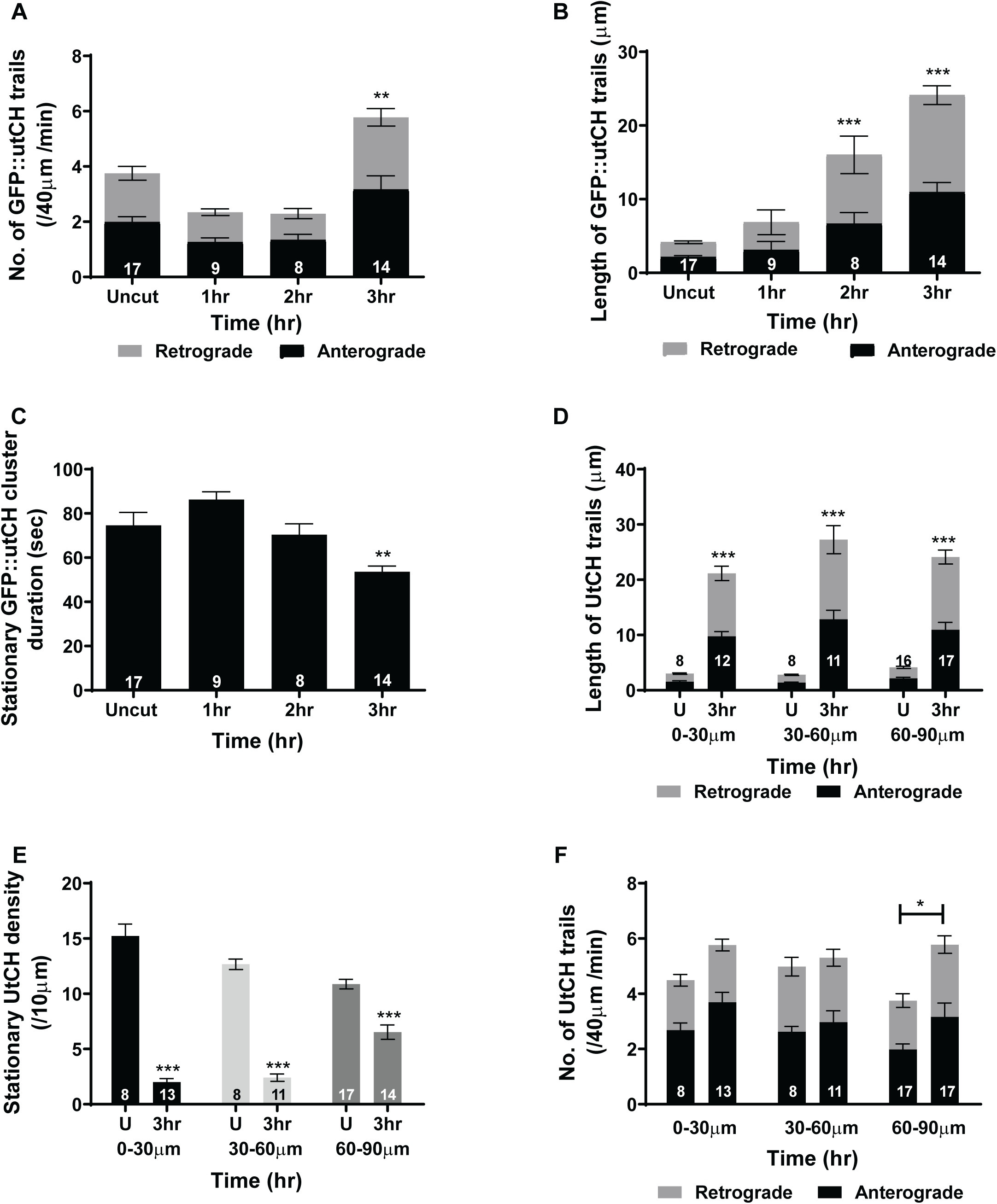
UNC-16 inhibits actin dynamics. ***A***. Number of GFP::utCH trails in neurons of uncut and axotomized wild type animals as a function of time up to 3 hr post injury in anterograde and retrograde direction. ***B***. Change in length of GFP::utCH trails in neurons of uncut and axotomized wild type animals over 3 hr post injury period in anterograde and retrograde direction. ***C***. Comparison of the numbers of stationary GFP::utCH stationary clusters in neurons of uncut and axotomized wild type animals over time period of 3 hr post injury. ***D***. Change in length of GFP::utCH trails in uncut and axotomized wild type animals at 3 hr post injury over the entire section of the neuron in anterograde and retrograde direction. ***E***. Comparison of the numbers of stationary GFP::utCH stationary clusters in uncut and axotomized wild type animals 3 hr post injury in various sections of the neuron. ***F***. Number of GFP::utCH trails in uncut and axotomized wild type 3 hr post injury over various sections of the neuron in anterograde and retrograde direction. Number of animals inside bars. One-way ANOVA p-value *<0.05, **<0.005, ***<0.001.

The punctate localization of *C. elegans* GFP::DLK-1S depends on its association with DLK-1L (Yan & Jin, 2012). GFP::DLK-1S, similar to GFP::DLK-1L is not punctate in *unc-16(lf)* animals (Sup. Fig. 6, 7). Surprisingly, DLK-1S is punctate upon over-expression of T7::UNC-16 in the absence of DLK-1L, i.e. in a *dlk-1(null)* background (Sup. Fig. 6, 7). This indicates that UNC-16 can scaffold DLK-1L and DLK-1S independent of each other, although our experiments cannot address whether it can scaffold a dimer of DLK-1L and 1S.

**Figure 6.**
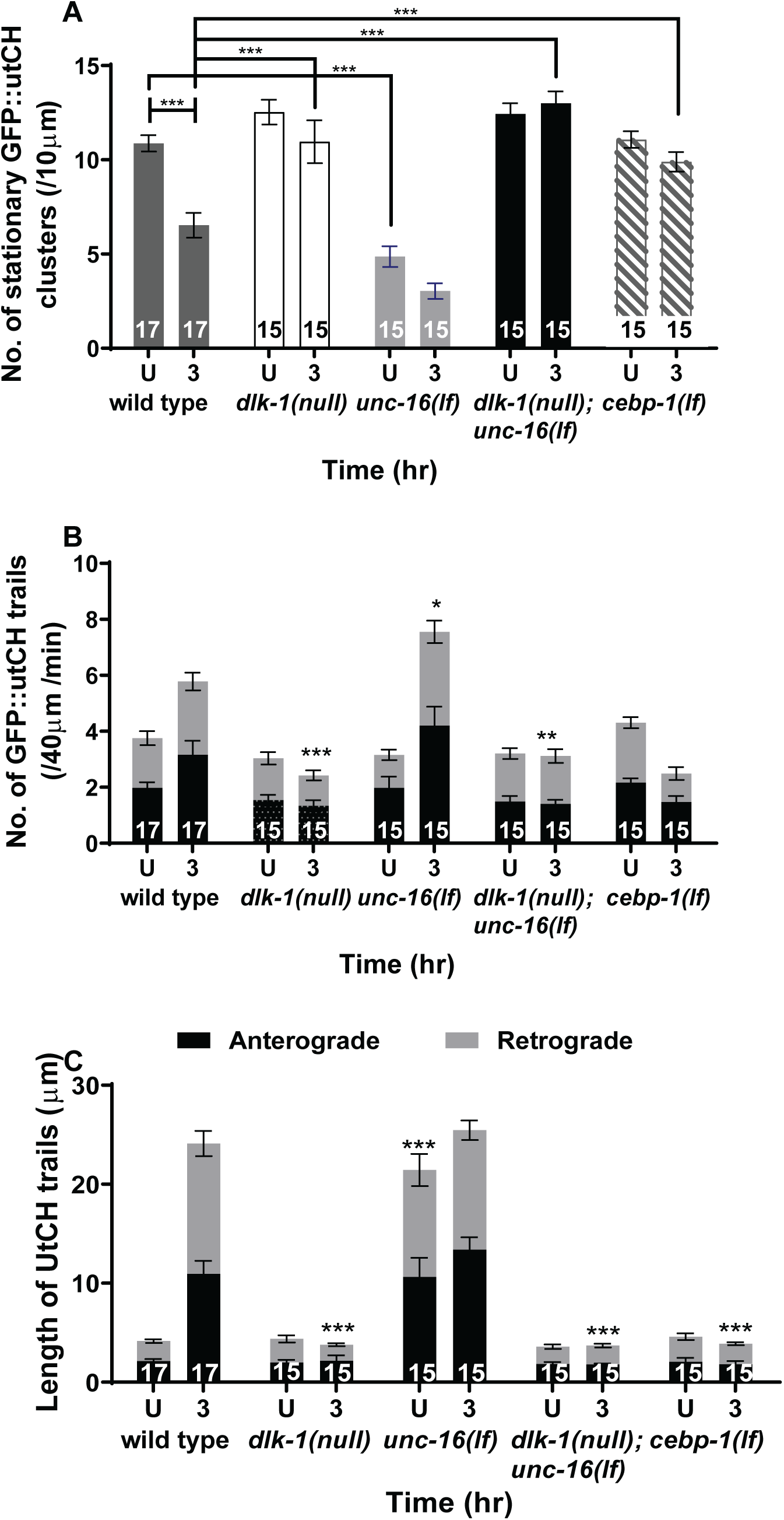
Changes in actin dynamics post-injury are dependent on DLK-1. ***A***. Comparison of the numbers of stationary GFP::utCH stationary clusters in uncut and 3 hr post injury neurons in various genotypes. ***B***. Number of GFP::utCH trails in uncut and 3 hr post injury neurons in various genotypes in anterograde and retrograde direction. ***C***. Comparison of the length of GFP::utCH trails in uncut and 3 hr post injury neurons in various genotypes in anterograde and retrograde direction. Number of animals inside bars. One-way ANOVA p-value *<0.05, **<0.005, ***<0.001.

Upon axotomy DLK-1L is known to increase locally at the injury site while the DLK-1S does not (Yan & Jin, 2012). We examined whether the post-axotomy localization of DLK-1L is dependent on UNC-16 and whether the sequestering of DLK-1L along the axon altered post-axotomy. Like the earlier report we observe that 4 minutes after axotomy, the levels of GFP::DLK-1L at the proximal end of the cut site increases (Fig. 3C, D). However, in *unc-16(lf)* there is a greater accumulation of GFP::DLK-1L at the proximal cut site (Fig. 3C, D) compared to wild type at the same time point. Further, overexpression of UNC-16 leads to a great reduction of GFP::DLK-1L accumulation at the proximal cut site with the GFP::DLK-1L signal comparable to the GFP alone control (Fig. 3C, D). Thus, the presence and levels of UNC-16 controls the post-axotomy levels of DLK-1L at the cut site. Additionally, the intensity of DLK-1L punctae along the neuronal process ∼10 um away from the injury site reduces (Fig. 3C). The extent of reduction in intensity depends on the presence and levels of UNC-16 (Fig. 3C).

The UNC-16 dependent localization of DLK-1L before injury and UNC-16 dependent control of DLK-1L levels post-injury at the cut site and along the neuronal process suggests that UNC-16 may act to sequester DLK-1L along the axon. The release of DLK-1L from the punctae may be stimulated by injury signals like Ca+2 known to bind DLK-1L (Yan & Jin, 2012). Taken together, our data suggest that UNC-16 may contribute to making DLK-1L unavailable at the cut site, perhaps by sequestering it in punctae along the neuronal process.

### DLK-1 controls both microtubule and actin dynamics

DLK-1 is known to enable neuronal regeneration through two independent pathways, by elevating microtubule dynamics and by regulating transcription through CEBP-1 (Ghosh-Roy, Wu, Goncharov, Jin, & Chisholm, 2010; Li, Hisamoto, & Matsumoto, 2015; Nix, Hisamoto, Matsumoto, & Bastiani, 2011; Tang & Chisholm, 2016; Yan et al., 2009) (Fig. 7D). Additionally actin, the actin depolymerising factor (ADF)/cofilin A and nonmuscle Myosin II that powers actin’s retrograde flow are needed for growth cone formation and for the extension of both developing and regenerating vertebrate neurons (Blanquie & Bradke, 2018; Chisholm, Hutter, & Jin, 2016; Flynn et al., 2012; Hur et al., 2011; Tedeschi et al., 2019). Thus we examined the dynamics of both microtubules and actin in wild type and in *dlk-1(0)* animals. Since we cannot assess depolymerisation events, the cytoskeletal dynamics that we report here are an evaluation of microtubule and actin polymerisation events.

**Figure 7.**
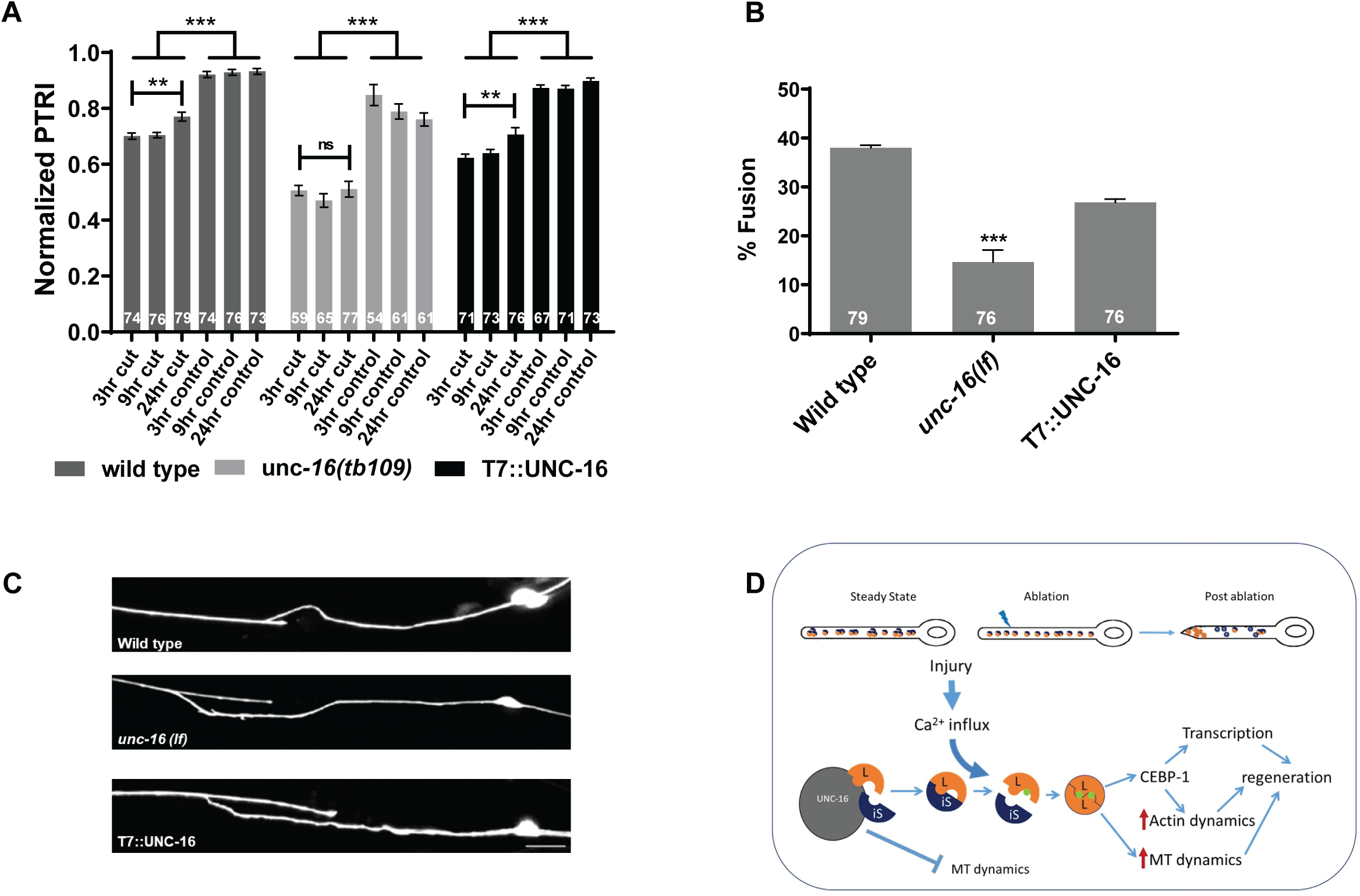
*unc-16(lf)* does not enhance functional recovery. ***A***. Posterior touch response index (PTRI) in axotomized and internal control neurons in wild type, *unc-16(lf)* and T7::UNC-16 animals. Multiple column comparison of ANOVA, with Neumann Keul’s post-test p-value *<0.01, **<0.001 and ***<0.0001. ns: not significant. ***B***. Percentage of animals showing neuronal fusion of the proximal and distal end post-axotomy at 9 hr in wild type, *unc-16(lf)* and T7::UNC-16 animals. One-way ANOVA p-value ***<0.001. Number of animals inside bars. ***C***. Representative images of neuronal fusion of the proximal and distal end post-axotomy at 9 hr in wild type, *unc-16(lf)* and T7::UNC-16 animals. Scale bar 10 µm. D. Suggested model of mechanism of inhibition of neuronal regeneration by UNC-16.

We assessed microtubule dynamics using EBP-2::GFP, a marker for the growing plus end of microtubules (Ghosh-Roy, Goncharov, Jin, & Chisholm, 2012). We measure the total number of EBP-2::GFP comets which are visualised as sloped lines (arrow) in kymographs (Sup. Fig. 8), length of the comets and direction of the comets (Sup. Fig. 9). In the wild type uncut animals, the number of EBP-2::GFP comets pre- and immediately post-axotomy in a 40 µm adjacent to the axotomy site are similar (Fig. 4A, B, Sup. Fig. 10). It has been shown that EBP-2 dynamics changes only in this region, remaining unchanged in the other regions of the proximal neuronal process post-axotomy (Ghosh-Roy et al., 2012).

We tracked microtubule dynamics up to 6 hours post-axotomy. The number of EBP-2::GFP comet tracks increases 3 hours post-axotomy (Fig. 4A, B). Three and six hours post-axotomy the number of EBP-2::GFP comets show a two-fold and 2.5-fold significant increase which are comparable to earlier reports (Ghosh-Roy et al., 2012). In wild type animals the length of the EBP-2::GFP comet tracks shows a peak increase 2 hours post-axotomy (Fig. 4C) that continues 3 hours post-injury with no further increase 6 hours post-injury. In uninjured *dlk-1(0)* neurons, the number of EBP-2::GFP comets are fewer than in wild type animals and did not significantly increase even 6 hours post-axotomy (Fig. 4A). We see a similar trend where EBP-2::GFP track length increases 3 hours post-axotomy but at 6 hours post-axotomy, there is no change compared to the uninjured neurons of *dlk-1(0)* animals.

We followed actin dynamics by using a transgenic line that expresses the GFP tagged calponin homology domain of F-actin-binding protein Utrophin (GFP::utCH) as a marker of F-actin-rich regions (Burkel, von Dassow, & Bement, 2007; Chia, Chen, Li, Rosen, & Shen, 2014; Sood et al., 2018). We measured three parameters, viz., the density of immobile actin-rich regions, number of actin trails and actin trail length as described earlier (Ganguly et al., 2015; Sood et al., 2018) (Sup. Fig. 9). Immobile actin-rich regions are visible in distance and time kymograph plots as vertical lines (arrow) and actin trails as sloped lines (arrowhead) (Sup. Fig. 11).

In the 40 µm proximal to the injury site, the number of GFP::utCH trails in the wild type animals increases significantly only 3 hours post-axotomy with an 1.6 fold increase in the anterograde direction and 1.4-fold increase in the retrograde direction (Fig. 5A). The length of the GFP::utCH trails increases three-fold, 2 hours post-axotomy in the anterograde direction and nearly 5 fold in the retrograde direction (Fig. 5B). In wild type animals the density of actin-rich regions reduce significantly 3 hrs post-axotomy (Fig. 5C). Unlike microtubule dynamics that show regional changes (Ghosh-Roy et al., 2012), actin dynamics changes throughout the proximal injured neuron 3 hours post-axotomy. We see an increase in the length of GFP::utCH trails in both the anterograde and retrograde directions (Fig. 5D) and a decrease in actin-rich regions throughout the axon (Fig. 5E). The changes are the greatest in the proximal 60 µm next to the axotomy site 3 hrs post-axotomy where the density of actin-rich regions reduces (Fig. 5D) and the length of anterograde and retrograde GFP::utCH trails increase the most (Fig. 5E) while the number of anterograde and retrograde GFP::utCH trails increase 60-90 µm away from the site of axotomy (Fig. 5F).

The density of actin-rich regions in uninjured *dlk-1(0)* mutants are similar in number to wild type (Fig. 6A). The numbers and lengths of anterograde and retrograde actin trails in wild type and *dlk-1(null)* are comparable as well (Fig. 6B, 6C). 3 hours post-axotomy there is no statistically significant change in density of actin-rich regions, numbers or lengths of actin trails in *dlk-1(null)* animals (Fig. 6). This shows that changes in actin dynamics post-injury are dependent on DLK-1.

Both microtubule and actin dynamics post-axotomy are DLK-1 dependent. However microtubule dynamics change only locally near the injury site (Ghosh-Roy et al., 2012) while actin dynamics changes throughout the axotomized neuronal process connected to the cell body (Fig. 5). This difference may arise from differential genetic control downstream of DLK-1 on actin and microtubules.

### UNC-16 regulates microtubule dynamics independent of DLK-1 and actin dynamics dependent on DLK-1

We assessed the effect of UNC-16 on microtubules and actin dynamics pre- and post-axotomy. We observe that there is an elevation in the number of EBP-2::GFP comets in uninjured neurons of *unc-16(lf)* (Fig. 4A). The double mutant *dlk-1(0); unc-16(lf)* continues to have an 1.5 fold significant increase in the number of EBP-2::GFP comets similar to the elevation in baseline microtubule dynamics seen in the *unc-16(lf)*. The number of EBP-2::GFP comets show nearly a two-fold increase 6 hr post-injury close to the 2.5 fold increase seen in wild type axotomized processes (Fig. 4A). The length of the EBP-2::GFP comet trails are elevated in wild type, *dlk-1(0)*, *unc-16(lf)* and *dlk-1(0);unc-16(lf)* three hours post-axotomy (Sup. Fig. 10). This upregulation of microtubule dynamics further increases post-injury in the neuron which is reflected in both the increased number and length of the EBP-2::GFP comets. In all the genotypes tested, there is no significant difference in the orientation of microtubule growth compared to the wild type (Sup. Fig. 12). Surprisingly although regeneration in *unc-16(lf)* is completely dependent on DLK-1 (Fig. 2A), the microtubule dynamics in *unc-16(lf)* is not dependent on DLK-1.

We then assessed the GFP::utCH dynamics in *unc-16(lf)* mutants. We observed that the length of the GFP::utCH trails is elevated five-fold in uninjured *unc-16(lf)* neurons compared to wild type (Fig. 6C). However, the number of GFP::utCH trails in uncut *unc-16(lf)* mutants is similar to that in wild type animals (Fig. 6B). The density of GFP::utCH marked actin-rich regions is lower by ∼50% in uninjured *unc-16(lf)* animals compared to wild type animals perhaps contributing to the polymerising F-actin pools in the neuronal process seen in this genotype (Fig. 6A). 3 hrs post-injury, the length of the GFP::utCH trails in *unc-16(lf)* mutants are similar to trails in axotomized wild type neurons (Fig. 6C). However, the numbers of trails are 1.3-fold greater in *unc-16(lf)* mutants compared to wild type animals 3 hours post-axotomy (Fig. 6B). In all the genotypes tested, the increase in GFP::utCH trails occurs in both anterograde and retrograde directions. Interestingly 3 hrs post-axotomy, *unc-16(lf)* mutants showed only a small non-significant reduction in density of actin-rich regions compared to uninjured mutants (Fig. 6A). By contrast in wild type there is a significant reduction in density of actin-rich regions post-injury. Double mutant *dlk-1(0);unc-16(lf)* was indistinguishable from *dlk-1(0)* and shows no increase in length, number and density of GFP::utCH trails in both uninjured neurons and 3 hrs post axotomy (Fig. 6) illustrating that the actin dynamics in *unc-16(lf)* is completely dependent on DLK-1.

We also investigated whether the altered actin dynamics changes the structure of the growth cone. Hence we counted the number of filipodial protrusions in wild type, *unc-16(lf)*, *dlk-1(0)* and *dlk-1(0);unc-16(lf)* animals. We observe that in *unc-16(lf)*, the genotype with greatly elevated actin dynamics pre- & post-injury, also shows an increase in the number of filipodial protrusions after injury compared to all remaining genotypes examined (Sup. Fig. 13, 14). This elevation in filipodial protrusions in *unc-16(lf)* begins as early as 6 hr (Sup. Fig. 15). Wild type animals show greater number of protrusions at the growth cone along with an elevation in actin dynamics post-injury by contrast to *dlk-1(0)* (Sup. Fig. 14). Thus, elevated actin dynamics pre- and post-axotomy may help in restructuring the growth cone and pre-program the neuron for swifter regeneration in *unc-16(lf)* animals. Actin dynamics likely make significant contributions to the elevated regeneration in *unc-16*, lack of regeneration in *dlk-1* and the lack of regeneration in *dlk-1; unc-16*.

In conclusion, cytoskeletal dynamics is elevated in *unc-16(lf)*. The elevated microtubule dynamics persists post-axotomy but is largely independent of DLK-1. By contrast, actin dynamics in *unc-16(lf)* is also elevated but is fully dependent on DLK-1. Since regeneration in *unc-16(lf)* is dependent on DLK-1 (Fig. 2A), regulation of actin dynamics via DLK-1 may be critical for the elevated regeneration seen in *unc-16(lf)* animals. However, the increased microtubule dynamics in *unc-16* may also play a DLK-1 independent role in regeneration, for instance in controlling the rate of axon outgrowth.

### UNC-16’s inhibition of actin dynamics is CEBP-1 dependent

DLK-1 is known to affect neuronal regeneration by promoting translation and stabilization of CEBP-1 mRNA in touch neurons (Yan et al., 2009) (Fig. 7D). CEBP-1 along with another transcription factor ETS-4 activates other growth factors needed for neuronal outgrowth during regeneration (Li et al., 2015). Additionally *cebp-1(lf)* animals do not show any change in the number of EBP-2 comets post-axotomy (Ghosh-Roy et al., 2012). Since the DLK-1 dependent regeneration in *unc-16* appears independent of microtubule dynamics, we assessed if the inhibitory role of *unc-16* on regeneration was mediated through the CEBP-1 pathway. Additionally, since actin dynamics changes throughout the neuronal process after axotomy, we also investigated whether this change was dependent on CEBP-1.

We assessed the effect of loss of UNC-16 in *cebp-1(0)* null mutants. We observed that despite complete absence of CEBP-1, the *unc-16(lf);cebp-1(0)* double mutants show a two-fold higher regeneration than *cebp-1(0)* alone 9 hr post-axotomy (p-value 0.04) (Sup. Fig. 16). This is unlike *dlk-1(0);unc-16(lf)* which shows no regeneration similar to the regeneration phenotype seen in *dlk-1(0)* (Fig. 2B). This data suggest that the regeneration observed in *unc-16(lf)* may not be entirely dependent on the CEBP-1 pathway.

We then examined whether actin dynamics is dependent on CEBP-1. Actin dynamics in *cebp-1(0)* does not change post-axotomy and is indistinguishable from *dlk-1(0)* pre- or post-axotomy (Fig. 6). We conclude that regulation of actin dynamics occurs through CEBP-1 suggesting that actin regulation can occur independent of changes in microtubule dynamics, potentially through CEBP-1’s action on regulators of actin.

### Functional recovery of the regenerating neuron

Functional regeneration post-axotomy has been observed in *C. elegans* motor and touch neurons (Abay et al., 2017; Basu et al., 2017; Ben-Yakar, Chronis, & Lu, 2009; Ding & Hammarlund, 2018). More specifically, the axonal fusion of the proximal and distal ends of touch neurons post-injury has been shown to be responsible for the observed functional regeneration in touch neurons (Abay et al., 2017; Basu et al., 2017). Since loss of UNC-16 causes faster regrowth we examined whether it led to quicker functional regeneration.

The bilaterally symmetric Posterior lateral microtubule neurons (PLM) mediate a gentle touch escape response. A worm that is moving backward responds to a gentle touch to its posterior side by reversing its direction of locomotion (Chalfie et al., 1985). This response can be quantitatively assayed and is represented as PTRI (posterior touch response index) (Basu et al., 2017). This touch assay was performed on both the sides (right and left) at 3, 9 and 24 hrs post axotomy where only one of the two PLM neurons was axotomized. The reversal response mediated by the side with the uninjured posterior touch neuron serves as an internal control for each animal whose behaviour was assessed post-axotomy.

After ascertaining that there is no difference between PTRI of uncut and laser mock cut animals in all the genotypes tested, the touch assay was performed 3, 9 and 24 hr post axotomy (Fig. 7A). The PTRI was normalised to the unaxotomised PTRI of each genotype (Basu et al., 2017). The morphology assessment of the axotomised PLM was done after 24 hr (Fig. 7B, C). In the wild type worms, 3 hr post-axotomy, there is a significant decrease of the touch response on the axotomized side compared to the response on the side of animal with uncut PLM neuron (Fig. 7A). This lowered PTRI recovered significantly 24 hr post injury and about 38% of the injured wild type PLM neurons show fusion between the proximal and distal processes (Fig. 7B, C).

In *unc-16(lf)* animals, upon axotomy, there is a significant decrease in touch response on the axotomized neuron side compared to the touch responsiveness on the uninjured side leading to a lower PTRI (Fig. 7A). However there is no significant increase in PTRI from 3 to 24 hr after injury, thus there is an absence of significant functional recovery after injury despite greater neuronal outgrowth (Fig. 7A). Concurrently we observe that only 14.5% of the injured neurons show fusion between the distal and proximal ends (Fig. 7B). In animals expressing T7::UNC-16 we found a significant decrease in PTRI 3 hrs after injury compared to the uncut worms of the same genotype (Fig. 7A). After 24 hr there is significant increase in PTRI from 3 hr and we observe about 26.3% fusion at this time point (Fig. 7B). Although the number of neurons that show fusion is slightly lower in T7::UNC-16, it is not statistically significant compared to wild type (p-value of 0.13), and the PTRI at 24 hr is comparable to wild type.

Taken together these data show that although there is faster growth in *unc-16(lf)* animals, it does not translate into elevated functional regeneration. In contrast in animals overexpressing UNC-16, though the growth rate is significantly reduced, the slower regrowth is observed to support significant functional recovery 24 hrs post-axotomy. Thus, rapid rates of regeneration in an injured axon may not necessarily lead to better functional recovery. Further, excessive sprouting in *unc-16(lf)* (Sup. Fig. 13, 14) may reduce fusion events important for functional recovery.

## Discussion

Our study shows that UNC-16 plays an inhibitory role on regeneration where its presence and levels act early to control both growth initiation and regrowth rate. These effects are dependent on the MAPKKK DLK-1, essential for regeneration in multiple models. There are few known inhibitors that act directly on DLK, the master regulator of regeneration (Asghari Adib, Smithson, & Collins, 2018; Chisholm, Hutter, Jin, & Wadsworth, 2016; Hammarlund et al., 2009; Valakh, Frey, Babetto, Walker, & DiAntonio, 2015). Our study suggests that UNC-16/JIP3 acts by sequestering DLK-1 along the neuronal process to control the levels of DLK-1 at the injury site. The regulation of DLK-1 levels activates a genetic cascade one of whose downstream effectors is altering actin dynamics throughout the neuron. This study shows the central role of DLK-1 dependent actin dynamics elevated through CEBP-1 as a key mediator of axonal regeneration. Our data suggest that in the absence of UNC-16, the neuron has a very dynamic cytoskeleton and is poised to grow immediately after injury. In the regeneration pathway UNC-16 acts as a clamp on cytoskeletal dynamics inhibiting both microtubule dynamics and actin dynamics. Intriguingly the rapid regrowth, at least partially mediated by altered cytoskeletal dynamics, does not help in functional recovery. Our study identifies a key role for UNC-16 in calibrating the rate of axon regrowth to enable functional recovery.

Systematic genetic screens in *C. elegans* identified that only ∼10% of genes screened inhibit neuronal regeneration while 83% of genes identified in the screen promote neuronal growth post-injury (Chen et al., 2011). UNC-16/JIP3 is a negative regulator that acts on DLK-1L independently and in addition to the inhibitory isoform DLK-1S (Fig. 3C). Since JIP3 is also known to form a complex with DLK (Holland et al., 2016; Kelkar et al., 2000), we propose that the effects of UNC-16 on DLK-1L might be direct. Apart from JIP3, DLK-1S is the only known inhibitor of DLK-1L (Yan & Jin, 2012). Although, the inhibitory DLK-1S like isoform has been suggested to be present in humans but its presence has not been verified in systems other than *C. elegans* (Sup. Table). Other identified negative regulators of regeneration like DIP-2 and NMAT-2 act through other parallel pathways and not directly on DLK-1 (K. W. Kim et al., 2018; Noblett et al., 2019). Thus, UNC-16 and its vertebrate homologue JIP3 may act as a general inhibitor of DLK mediated signalling in multiple species by altering its localisation in an injury dependent manner. UNC-16 may merely function to sequester rather than regulate DLK-1 kinase activity or in addition to sequestering DLK-1 may also inhibit its kinase activity. Though DLK-1 exists in a heterodimer form, we do not have an estimate of the proportion of the heterodimer *in vivo* since UNC-16 can also sequester DLK-1S in puncta independent of DLK-1L (Sup. Fig. 6, 7). Upon injury UNC-16 appears to release DLK-1L from punctate structures that accumulates at the cut site (Fig. 3C, D). This free DLK-1 activates downstream pathways that ultimately lead to increased neuronal regeneration (Fig. 7D) (Yan & Jin, 2012). The sequestration of DLK may depend on palmitolyation (Holland et al., 2016) and thus UNC-16 bound DLK is likely to be palmitoylated and additional steps including removal of the palmitoyl linkage may control the levels of free DLK-1. Other members of the JIP family may also share a similar inhibitory roles on regeneration since JIP1 is also known to both directly bind to as well inhibit DLK activity (Nihalani, Meyer, Pajni, & Holzman, 2001).

Cytoskeletal damage is thought to trigger DLK activity that subsequently initiates regeneration post injury (Valakh et al., 2015). Our data suggests that the initiation of regrowth may require both microtubule and actin dynamics and both dynamics change at similar time scales after injury (Fig. 4, 5, 6). DLK-1 regulates local microtubule dynamics close to the injury site through the depolymerizing kinesin-13 family member, KLP-7 (Ghosh-Roy et al., 2012). Thus some of the elevated microtubule dynamics observed in *unc-16* may be dependent on KLP-7. Additionally, *unc-16* also shows elevated microtubule dynamics independent of DLK-1 (Fig. 4A, Sup. Fig. 11). However the dependence of elevated microtubule dynamics in *unc-16* on KLP-7 remains to be investigated. Another motor, Kinesin-1, is shown to be regulated by mammalian JIP3 to promote axon elongation (Sato et al., 2015). However *unc-16’*s effect on regeneration in *C. elegans* touch neurons appears independent of Kinesin-1 (Fig. 2A). Additionally, DLK-1 action on microtubule through molecular motors is likely to be local since the changes in microtubule dynamics have been shown to occur largely near cut site (Ghosh-Roy et al., 2012).

Our data suggest that UNC-16 controls actin dynamics through CEBP-1 (Fig. 6). Actin is thought to be important in the regenerating growth cone (Blanquie & Bradke, 2018; Chisholm, Hutter, & Jin, 2016; Flynn et al., 2012; Hur et al., 2011; Tedeschi et al., 2019). Overexpression of the actin depolymerising factor (ADF)/cofilin has been shown to induce regeneration in culture as well as *in vivo* following spinal cord injury (Tedeschi et al., 2019). Inhibition of nonmuscle myosin II, which powers retrograde actin flow in the growth cone, markedly enhances axon growth by reorganizing the growth cone cytoskeleton in cultured vertebrate neurons (Hur et al., 2011). The altered actin dynamics seen in *unc-16* may occur through molecules such as Cofilin, Myosin II and actin polymerizers such as Formin and ARP2/3 (Blanquie & Bradke, 2018; Breitsprecher & Goode, 2013; Chisholm, Hutter, & Jin, 2016; Flynn et al., 2012; Hur et al., 2011; Tedeschi et al., 2019). The transcription of some of these genes may be elevated by CEBP-1 post-injury in a DLK-1 dependent manner. Perhaps this transcriptional control regulates altered actin dynamics in the entire length of the injured neuron (Fig. 5D, F).

Our data suggests that dynamic actin is important in enabling regeneration post-injury. *unc-16(lf)* mutants have elevated dynamics of both microtubule and actin which may pre-program these mutant neurons for a head start in their regrowth response post-injury. In *unc-16*, the pre-existing greater dynamic actin may be responsible for the growth cone with greater protrusions and earlier initiation of regrowth while the elevated microtubule dynamics may also be responsible a faster long-term regrowth of the regenerating neuron. By contrast, the wild type has a delay in actin and microtubule dynamics post-injury leading to a slower initiation and slower growth rate at which the injured neuron regrows (Fig. 1D, E). However, this head start by *unc-16* is lost and wild type growth catches up in the later time points such that at 24 hour post-axotomy *unc-16(lf)* shows comparable axon extension (∼27 µm) to wild type (∼36 µm) regenerating neurons (Fig. 1D). DLK-1 dependent microtubule dynamics alone may be insufficient for neuronal regeneration post-axotomy since in *unc-16; dlk-1* animals microtubule dynamics is elevated but animals show no regeneration (Fig. 2A, 4A). A combination of both actin and microtubule dynamics may be important for successful neuronal regeneration. Despite differences in pathways downstream of DLK-1 in regulating both microtubule and actin dynamics, regrowth after injury is likely to depend on the interaction between the actin and microtubule cytoskeletons during growth cone formation (Dogterom & Koenderink, 2018). We propose a model where UNC-16/JIP3 plays its inhibitory role through tight temporal and spatial control of DLK-1 function (Fig. 7D). The dual inhibitory control by both UNC-16 and DLK-1S calibrate the intrinsic growth promoting function of DLK-1L *in vivo*. This intrinsic growth ability is likely achieved through a balance between the actin dynamics and microtubule stabilisation at the growth cone that could also mediate rapid steering of the growth cone in a regenerating neuron (Blanquie & Bradke, 2018; Hur et al., 2012).

JIP3 could play a dual role post-injury, through its effects on microtubule and actin dynamics as well as through its role in retrograde transport. UNC-16 and its mammalian orthologue JIP3 have been shown to associate with the retrograde motor dynein (Arimoto et al., 2011; Cavalli, Kujala, Klumperman, & Goldstein, 2005; Holland et al., 2016; Koushika et al., 2004; Terenzio et al., 2018). In injured neurons dynein transports importins, importin associated proteins, transcription factors and cargoes that can bind scaffolding molecules (Mahar & Cavalli, 2018; Rishal & Fainzilber, 2014). Spatial translocation of MAP kinases like Erk-1 and Erk-2 by importins and dynein is also reported in lesioned nerve and is shown to be important for regeneration (Hanz et al., 2003; Perlson et al., 2005; C. L. Wu et al., 2012). The dependence of movement of these cargoes on JIP3/UNC-16 is not yet investigated. Although our experiments suggest that the number of animals that show regenerative growth in *unc-16* is independent of dynein (Sup. Fig. 3), a dynein-dependent role in regulating the rate of neuronal outgrowth, a prerequisite to achieve functional regeneration, has not been examined.

We show that the faster regeneration seen in *unc-16(lf)* mutants does not lead to functional recovery known to depend on fusion of the distal and proximal parts of the injured neuron (Abay et al., 2017; Basu et al., 2017). The quicker and greater neuronal growth observed in *unc-16(lf)* mutants may not be optimal for fusion as it may adversely affect other required steps in a fully functional neuron, e.g., cargo transport that is known to depend on UNC-16/JIP3. The *efa-6* mutant that shows faster regrowth and elevated microtubule dynamics also does not show enhanced functional recovery (Basu et al., 2017; Chen et al., 2015). Further, slower regeneration seen when UNC-16 is over-expressed shows functional recovery comparable to wild type animals (Fig. 1D, 7A). Thus merely a faster rate of neuronal outgrowth is insufficient for functional recovery highlighting the importance of mechanisms that modulate the growth rate of the regenerating neuron to allow for recovery of function.

## Materials and Methods

### Strains

*C. elegans* strains were grown on NGM agar plates seeded with *E. coli* OP50 bacteria at 16°C (Brenner, 1974) (Table 1). *jkk-1p::T7::unc-16* plasmid was made by inserting T7-UNC-16 DNA excised from CMVT7p-UNC-16 plasmid (Byrd et al., 2001) into NHjkkp-1p, which was made by replacing *jkk-1p*::*gfp* DNA region of MK104p (Kawasaki et al., 1999) with multiple cloning sites. *ofm-1p*::DsRed-monomer has been described previously (Arimoto et al., 2011). The homozygosity of all MosSCI* (Single copy insertion of a transgene into a defined site) strains was confirmed using PCR before and after completion of the axotomy assays using reported primers (Yan & Jin, 2012).

**TABLE 1.**
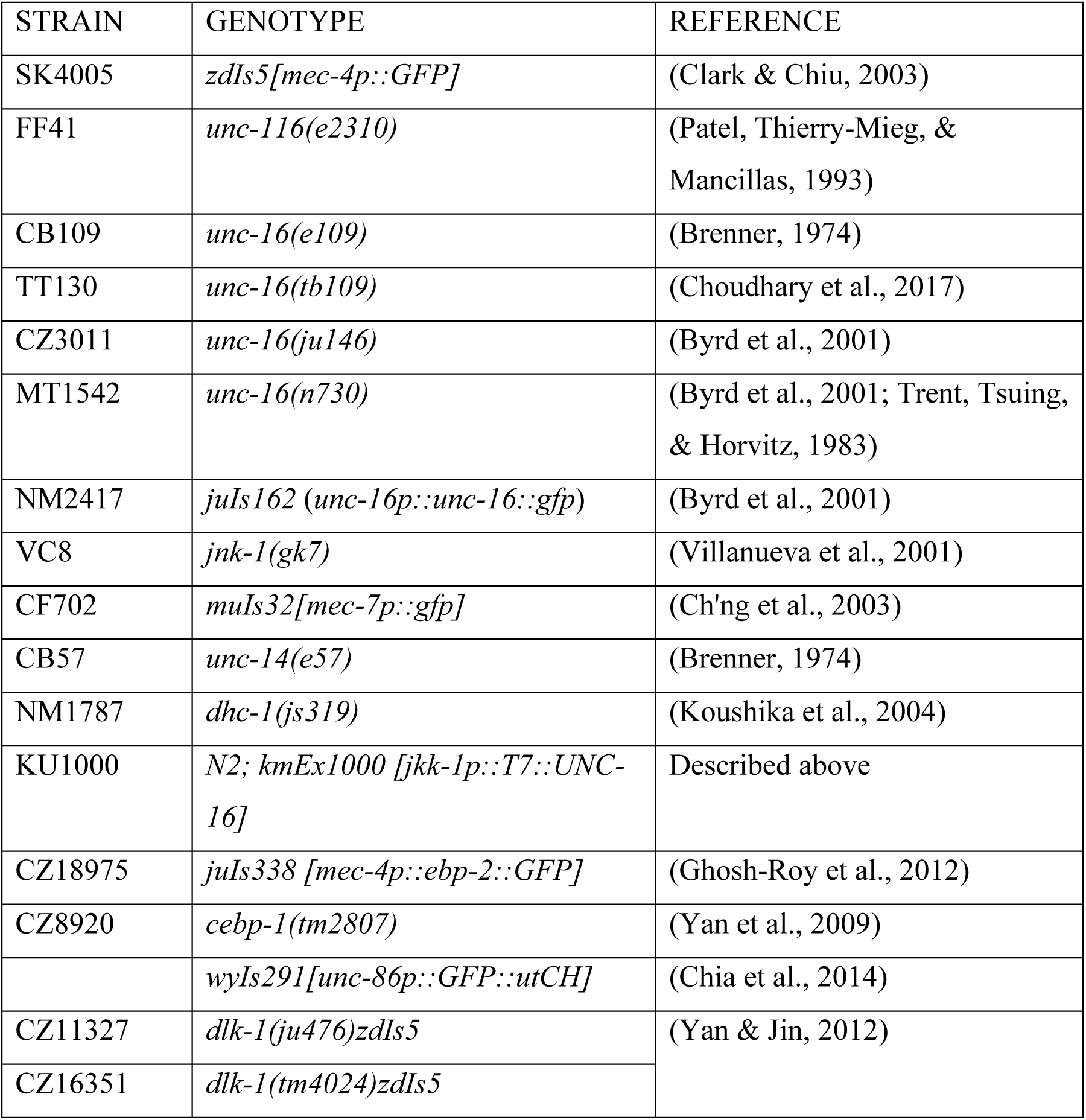
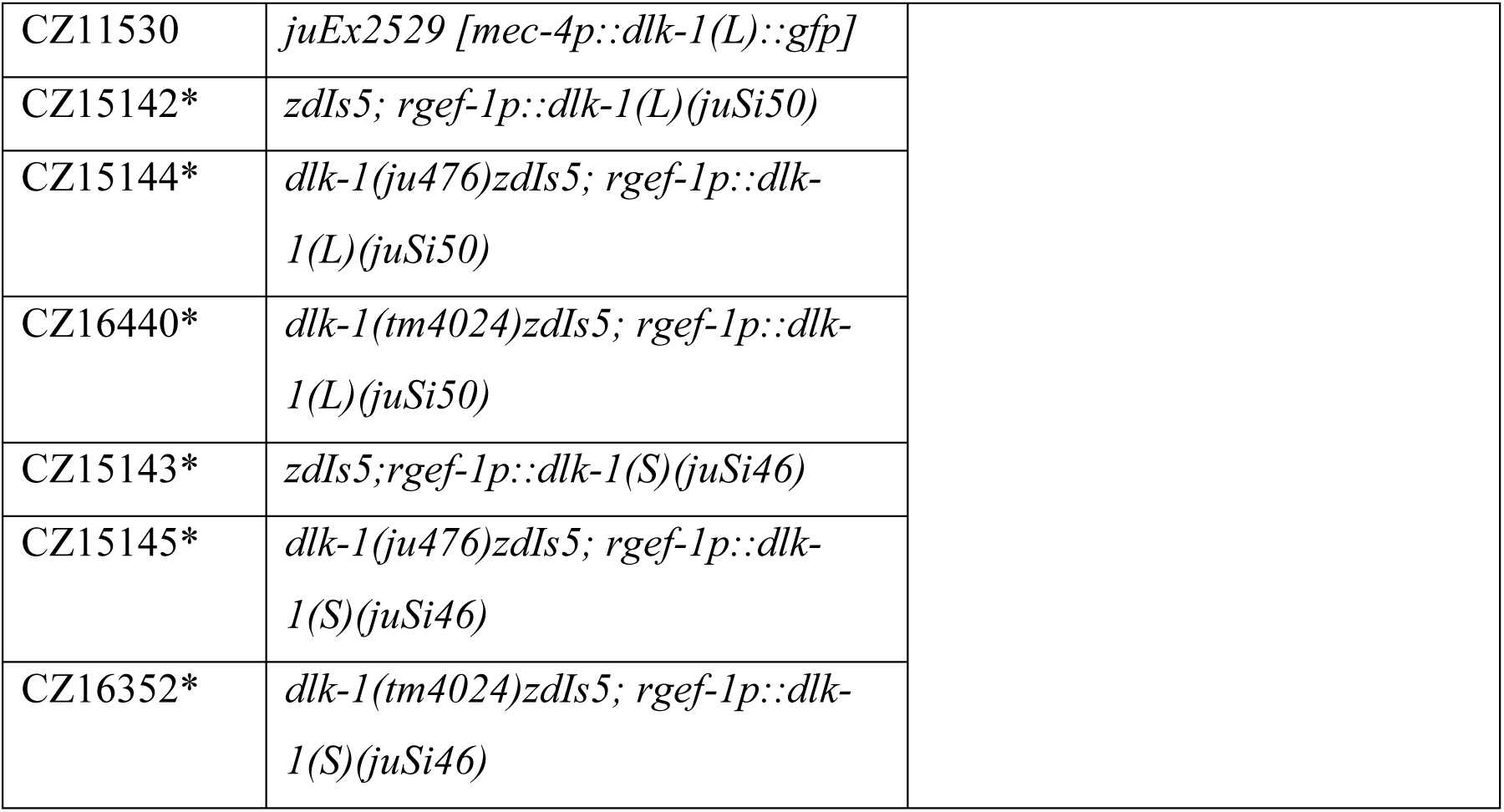
List of *C. elegans* strains

### Laser Axotomy

Nanosecond laser (Spitlight 600, Innolas, Munich, Germany) pulses (λ = 355 nm) of energy 0.8 μJ in single shot mode was used to generate neuronal cuts (Rao, Kulkarni, Koushika, & Rau, 2008). One of the Posterior Lateral Mechanosensory (PLM) neurons expressing soluble GFP was cut approximately 30-50 μm away from the cell body. All experiments were done using larval stage 4 animals grown at 16°C. We chose this temperature as the *unc16(tb109)* animals show accelerated growth and we wanted to ensure that we capture the characteristics of neuronal regeneration in *unc-16(tb109)* vis-à-vis wild type.

### Imaging and analysis

Images for analysis were acquired using a 100X objective, 1.4 NA at 1×1 binning and were processed using ImageJ software. Regeneration was represented as percent neuronal regeneration i.e. number of neuronal processes showing regrowth out of total number of axotomised neurons (Hammarlund et al., 2009; Nix et al., 2011). The length of the regrowing process was measured from the cut site (Z. Wu et al., 2007; Zou et al., 2013). Neurons showing >4 µm regrowth were considered as regenerating. Imaging was done 9 hours post-axotomy unless otherwise mentioned.

### Statistical Analysis

The data has been represented as mean ± s.e.m. The data when represented as % regeneration is an average of at least two separate trials. One-way ANOVA was used as a test of significance unless specified otherwise. p-values<0.05 were considered as significant.

### Localization of GFP tagged DLK isoforms

PLM neurons of one-day adult animals were imaged using a Perkin Elmer spinning-disc confocal microscope (488 nm laser, objective 100X, 1.4 NA) with a EMCCD C9100-13 camera.

### Microtubule and actin dynamics measurement

Laser axotomy was performed using a ns-pulsed Nd-YAG LASER at 355 nm and energy 0.8 mJ (Litron Nano S60-30) using a 60X, 1.35 NA objective. The axotomy was performed on *juIs338* or *wyIs291* L4 animals 100 μm from the cell body (Sup. Fig. 9). The animals were washed with M9 buffer and rescued. At the reported time point, the animals were anesthetized with 5 mM Levamisole and imaged using 488 nm laser excitation with a Yokogawa CSU-X1-A3 spinning disc confocal head and a Hamamatsu C9100-13 EMCCD Camera controlled by Volocity collectively supplied by PerkinElmer. Imaging was done under 100X 1.4 NA objective using 300 ms exposure at 3 frames/second for a total of 3 minutes for *juIs338* and 600 ms exposure at 1.6 frames/second for a total of 3 minutes for *wyIs291*. Kymographs were generated using the ImageJ plugin MultiKymograph (http://imagej.net/Multi_Kymograph). EBP-2::GFP comets and GFP::utCH trails with polymerizing length above 0.3 mm were used for total number, speed and length of EBP-2::GFP comet. The total number of EBP-2::GFP comets and GFP::utCH trails were then normalized to the duration of the image file as well as the length of the ROI in focus to generate the normalized microtubule dynamics graph.

### Behavioural assay

Laser axotomy was performed using a multiphoton IR laser (720 nm) on one PLM 50 µm away from the cell body. Two cuts were introduced 6 µm apart. A worm that is moving forward responds to a gentle touch in the anterior side by reversing, and a worm that is moving backwards likewise responds to a posterior touch by reversing. This touch assay was performed after 3, 9 and 24 hrs post axotomy. The phenotyping of the axotomised PLM was done after 24 hrs. Using this assay, one can measure touch sensation with a posterior touch response index (PTRI) (Basu et al., 2017) which has been plotted. For statistics, multiple column comparison of ANOVA, with Neumann Keul’s post-test was used. *p<0.01, ** p<0.001 and *** p<0.0001.

## Acknowledgements

We thank Y. Jin and K. Miller for strains. CGC provided strains and is funded by NIH Office of Research Infrastructure Programs (P40 OD010440). We thank H. Krishnamurthy, M. Mathew at CIFF, NCBS. We thank M. Kamak for illustrations, A. Ghose and G. Hasan for suggestions.

## Author contributions

S.S.K., V.S., S.S. and S.P.K. designed experiments and wrote the manuscript. S.S.K., V.S. S.S., A.B. performed experiments, K.M., N.H. and A.G-R. provided reagents and methodology. All authors read the manuscript.

---

**Supplemental Figure 1.** Results of quantitative RNA estimation in various alleles of *unc-16*. One-way ANOVA, p-value *<0.05.

**Supplemental Figure 2.** Neuronal regeneration in *unc-116(e2310)* and *jnk-1(gk7)* and their doubles with *unc-16(tb109).* One-way ANOVA p-value *<0.05, ***<0.001. Number of animals inside bars.

**Supplemental Figure 3.** Neuronal regeneration in *unc-14(e57)*, *dhc-1(js319)* and *dlk-1(ju476)* and their doubles with *unc-16(tb109)*. One-way ANOVA p-value *<0.05, ***<0.001. Number of animals inside bars.

**Supplemental Figure 4.** Schematic of DLK-1L and DLK-1S.

**Supplemental Figure 5.** Puncta density quantification of DLK-1L::GFP in *unc-16(tb109)* and T7::UNC-16 and other alleles of *unc-16*. One-way ANOVA p-value *<0.05, **<0.005, ***<0.001. Number of animals inside bars.

**Supplemental Figure 6.** Normalised intensity quantification of effect of *unc-16(tb109)* and T7::UNC-16 on GFP::DLK-1S localization. Intensity of *mec-4p::mCherry* was used as a negative control. One-way ANOVA p-value **<0.005, ***<0.001. Number of animals inside bars.

**Supplemental Figure 7.** Representative picture of GFP::DLK-1S localisation. Scale bar 10 µm.

**Supplemental Figure 8.** EBP-2::GFP track kymographs (inverted greyscale) observed in uncut and 3 hr post-injury neurons in wild type, *dlk-1(null)*, *unc-16(lf)* and *dlk-1(null);unc-16(lf)*. The length (x axis) and time (y axis) scales are 10 μm and 25 s, respectively.

**Supplemental Figure 9.** Schematic of the area of ablation and recording in the cytoskeletal studies.

**Supplemental Figure 10.** Length (μm) of EBP-2::GFP tracks in kymographs, before and after axotomy. One-way ANOVA p-value *<0.05, **<0.005, ***<0.001. Number of animals inside bars.

**Supplemental Figure 11.** Representative kymographs (inverted grayscale) of GFP::utCH trails prior to injury and 3 hr post injury in PLM neuron. The length (x axis) and time (y axis) scales are 10 μm and 25 s, respectively.

**Supplemental Figure 12.** Orientation of EBP-2::GFP tracks in kymographs (anterograde or retrograde), before and after axotomy in wild type and specified mutants. Number of animals inside bars.

**Supplemental Figure 13.** Representative pictures of the growth cone protrusions at different time points post injury in wild type and *unc-16(lf)* mutant neurons. Scale bar 5 µm.

**Supplemental Figure 14.** Number of protrusions at 9 hr post axotomy in the neurons of wild type, *unc-16(lf)*, *dlk-1(null)* and *cebp-1(null)* animals. One-way ANOVA p-value ***<0.001. Number of animals inside bars.

**Supplemental Figure 15.** Number of protrusions at specified time points post axotomy in the neurons of wild type and *unc-16(lf)* animals. Comparison at similar time point with One-way ANOVA p-value *<0.05, ***<0.001. Number of animals is 20 at each time point of each genotype except *unc-16(lf)* 24 hr, n=11.

**Supplemental Figure 16.** Neuronal regeneration in wild type, *unc-16(lf)*, *cebp-1(null)* and its double with *unc-16(lf).* One-way ANOVA p-value *<0.05, **<0.005, ***<0.001. Number of animals inside bars.

**Supplemental Table.** NCBI Accession numbers, classification based on length after Leucine Zipper Domain. *Ca2+ binding consensus sequence SDGLSD.

