## Supplementary Data for "UNC-16/JIP3 negatively regulates actin dynamics dependent on DLK-1 and microtubule dynamics independent of DLK-1 in regenerating neurons"

### Supplementary Figures

1

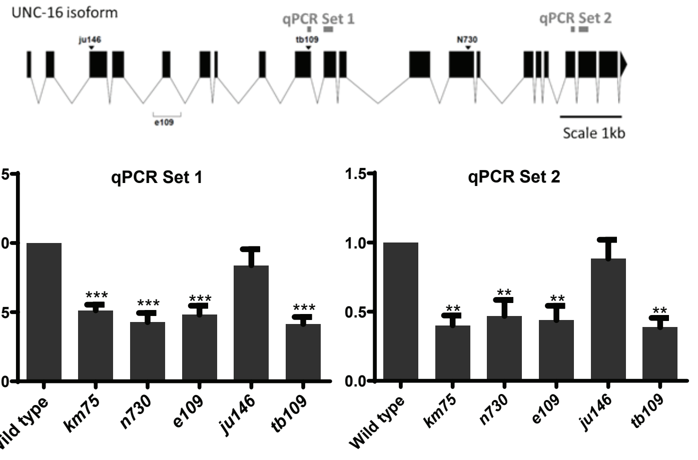


2

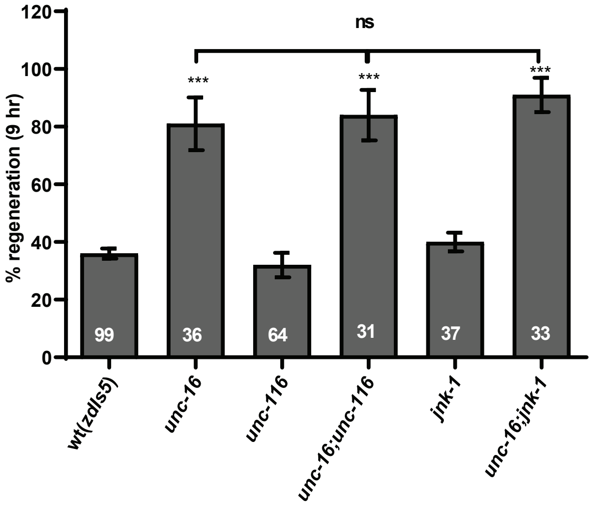


3

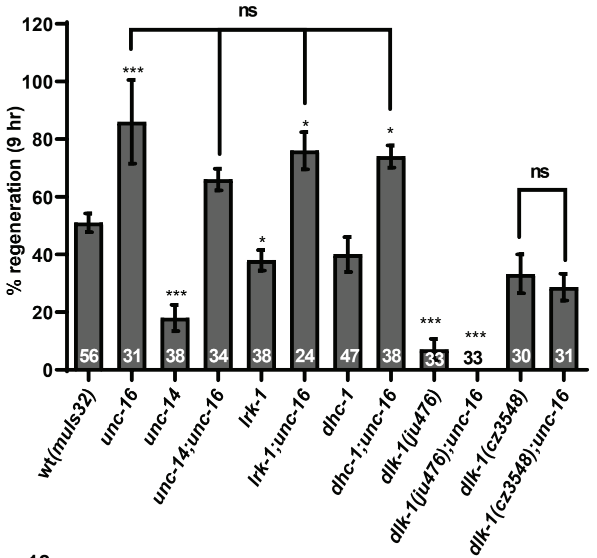


4

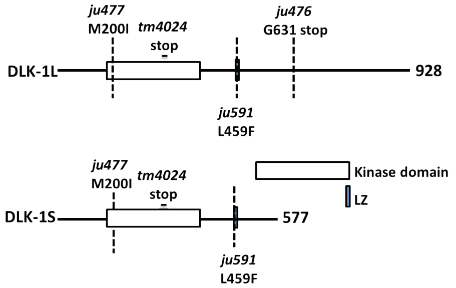


5

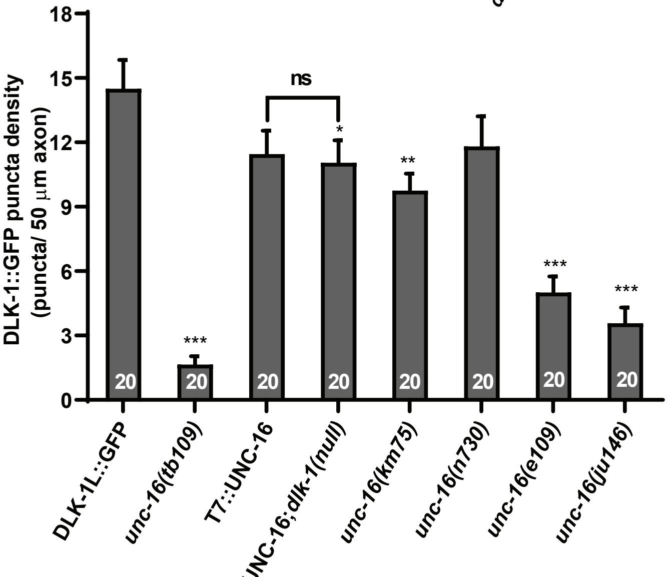


6

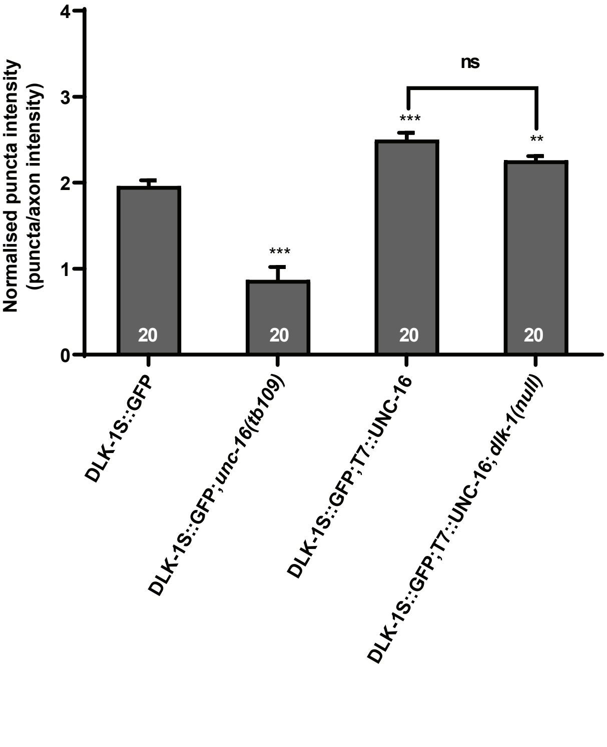

### Supplementary Figures

7

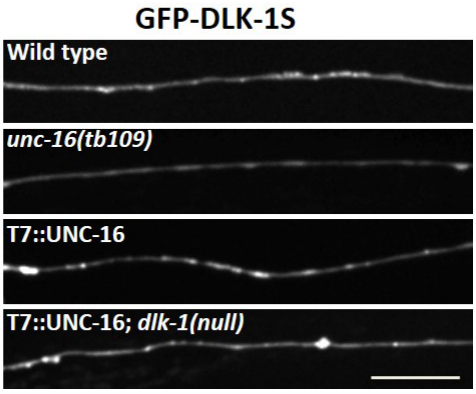


8

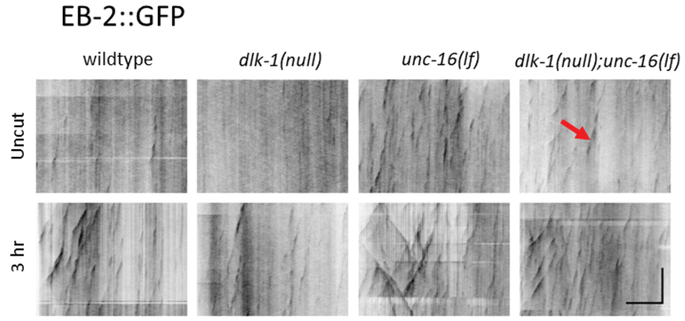


9

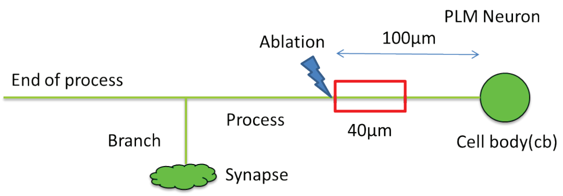


10

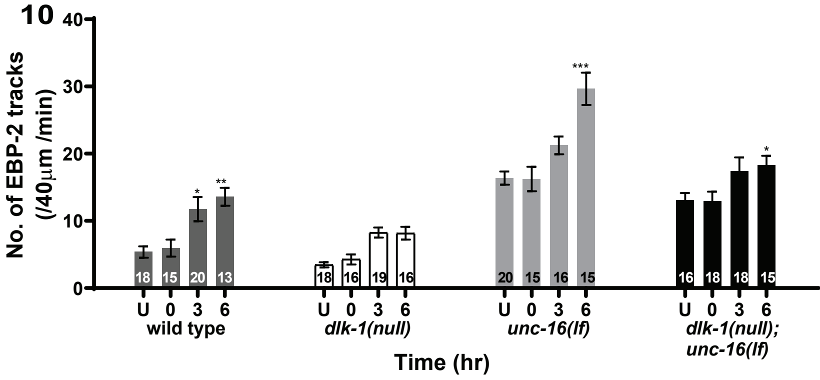


11

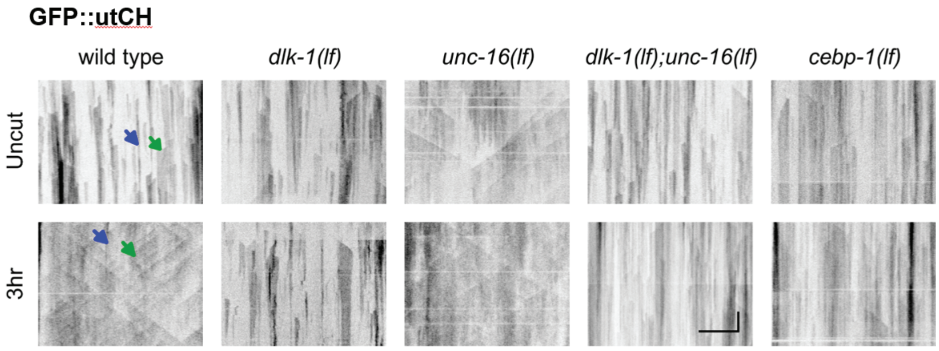

### Supplementary Figures

12

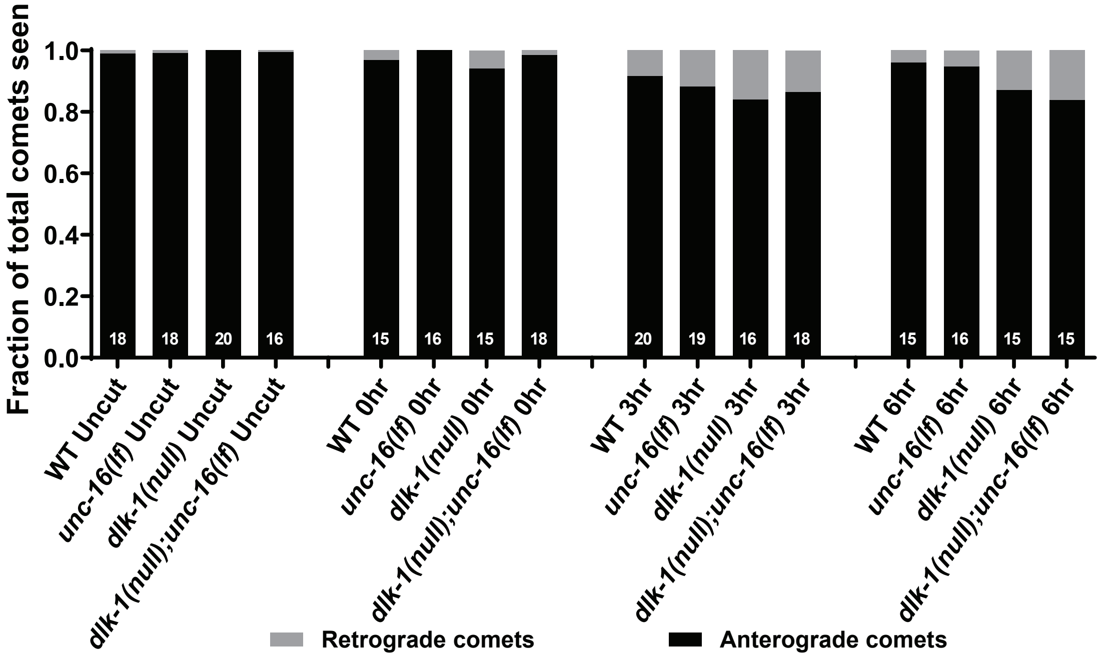


13

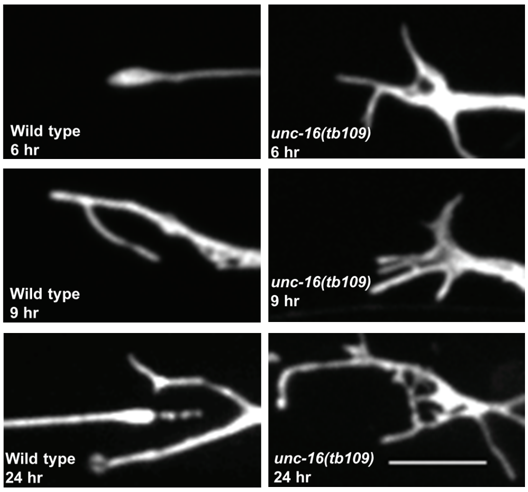


14

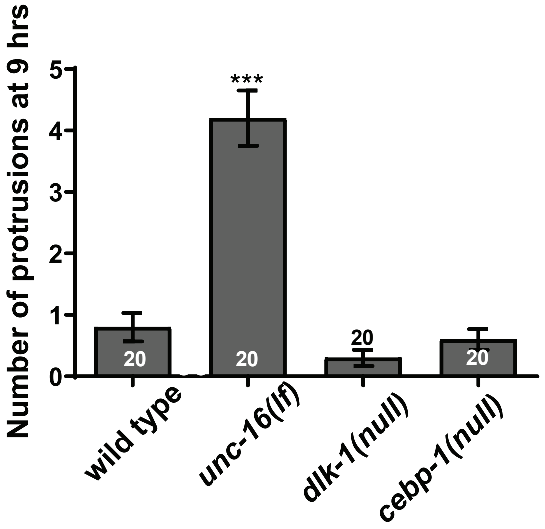


15

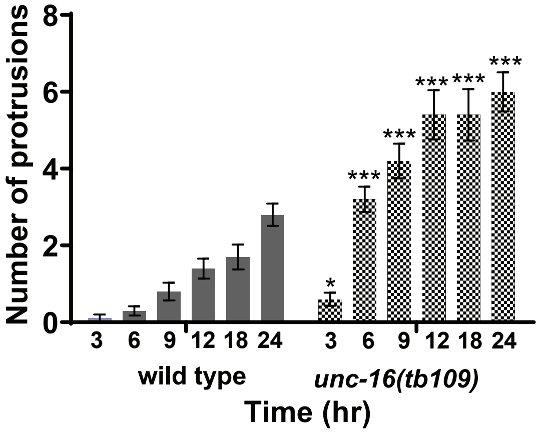


16

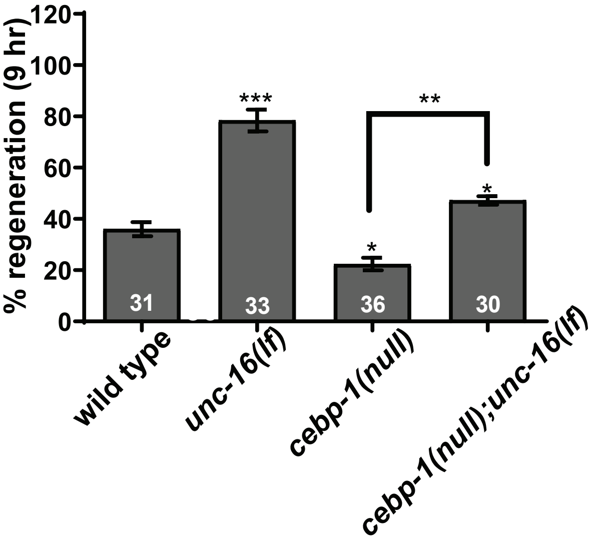

### Supplementary Table

| | **NCBI Accession numbers, classification based on length of Leucine Zipper Domain** | | | | | | |
|---|---|---|---|---|---|---|---|
| | *C. elegans* | *H. sapiens* | *M. musculus* | *R. norvegicus* | *B. taurus* | *D. rerio* | *D. melanogaster* |
| DLK-1 long like isoform | **DLK-1**<br>NP_001021444.1*<br>NP_001021443.1*<br>NP_001129769.1*<br>NP_001251937.1* | **MAP3K12**<br>NP_001180110.1<br>NP_00629.3<br>**MAP3K13**<br>NP_001229243.1*<br>NP_001229246.1*<br>NP_004712.1* | **MAP3K12**<br>NP_001157115.1<br>NP-033608.3<br>**MAP3K13**<br>NP_766409.2* | **MAP3K12**<br>NP_037187.1<br>**MAP3K13**<br>NP_001014000.2* | **MAP3K12**<br>NP_001192220.1<br>**MAP3K13**<br>NP_001095323.1* | **MAP3K12**<br>NM_207094.1<br>**MAP3K13**<br>E7FH13* | **Wallenda**<br>NP_788541.1 |
| DLK-1 short like isoform | **DLK-1**<br>NP_001021445.1 | **MAP3K12**<br>XP_005269197.1 | | | | | |

NCBI Accession numbers, classification based on length after Leucine Zipper Domain
*$Ca^{2+}$ binding consensus sequence SDGLSD
